# Cryo-EM structure of endogenous *Plasmodium falciparum* Pfs230 and Pfs48/45 fertilization complex

**DOI:** 10.1101/2025.02.13.638205

**Authors:** Melanie H. Dietrich, Jill Chmielewski, Li-Jin Chan, Li Lynn Tan, Amy Adair, Frankie M. T. Lyons, Mikha Gabriela, Sash Lopaticki, Toby A Dite, Laura F Dagley, Lucia Pazzagli, Priya Gupta, Mohd Kamil, Ashley M. Vaughan, Rattanaporn Rojrung, Anju Abraham, Ramin Mazhari, Rhea J. Longley, Kathleen Zeglinski, Quentin Gouil, Ivo Mueller, Stewart A. Fabb, Rekha Shandre-Mugan, Colin W. Pouton, Alisa Glukhova, Shabih Shakeel, Wai-Hong Tham

**Affiliations:** The Walter and Eliza Hall Institute of Medical Research, Parkville, Victoria 3052, Australia; Department of Medical Biology, The University of Melbourne, Melbourne, Victoria 3010, Australia; Department of Infectious Diseases, Doherty Institute, University of Melbourne, Parkville, Victoria 3010, Australia; Seattle Children’s Research Institute, Seattle, WA, USA; Department of Pediatrics, University of Washington, Seattle, WA, USA; Division of Malaria Research, Proteo-Science Center, Ehime University, Japan; Faculty of Tropical Medicine, Mahidol University, Bangkok 10400, Thailand; Olivia Newton-John Cancer Research Institute, Heidelberg, Victoria 3084, Australia; School of Cancer Medicine, La Trobe University, Bundoora, Victoria 3086, Australia; School of Global Health, Shanghai Jiao Tong University, Shanghai 200025, China; Monash Institute of Pharmaceutical Sciences, Monash University, Parkville, Victoria 3052, Australia; Department of Biochemistry and Pharmacology, The University of Melbourne, Melbourne, Victoria 3010, Australia; Drug Discovery Biology, Monash Institute of Pharmaceutical Sciences, Monash University, 381 Royal Parade, Parkville, Victoria 3052, Australia; ARC Centre for Cryo-electron Microscopy of Membrane Proteins, Monash Institute of Pharmaceutical Sciences, Monash University, 381 Royal Parade, Parkville, Victoria 3052, Australia; Research School of Biology, The Australian National University, Canberra, ACT 2600, Australia

**Keywords:** malaria, vaccine, transmission blocking, Pfs230, Pfs48/45, fertilization, cryo-EM, nanobody, mRNA-LNP, *Plasmodium falciparum*

## Abstract

*Plasmodium falciparum* Pfs230 and Pfs48/45, part of a core fertilization complex, are leading malaria transmission-blocking vaccine candidates. However, how the two proteins interact is unknown. Here we report a 3.36 Å resolution cryo-electron microscopy structure of the endogenous Pfs230-Pfs48/45 complex. We show that Pfs48/45 interacts with Pfs230 domains 13 and 14, domains that are not included in current Pfs230 vaccine immunogens. Using a transgenic parasite line with a domain 13 to 14 deletion, we show that these domains are essential for Pfs230 localization on the gamete surface. Furthermore, this line significantly reduced oocyst formation in the mosquito midgut, showing that the presence of Pfs230 domains 13 and 14 is critical for successful fertilization. Nanobodies against domains 13 and 14 inhibit Pfs230-Pfs48/45 complex formation, reduce transmission and structural analyses reveal their binding epitopes. Furthermore, domains 13 and 14 are targets of naturally acquired immunity and when delivered as mRNA-LNP immunizations induce potent immune responses and blocked transmission of malaria parasites. Our comprehensive structural insights on a core *P. falciparum* fertilization complex will guide the design of novel transmission-blocking vaccine candidates against malaria.

## Main text

Malaria is a major disease in humans causing over 600,000 deaths each year, with *Plasmodium falciparum* being responsible for almost all deaths^1^. *P. falciparum* has a complex life cycle involving the female *Anopheles* mosquito and a vertebrate host, the human. Development of highly effective vaccines against malaria has been hindered by the complexity of the parasite life cycle, with the parasite undergoing vast morphological and antigenic changes across its pre-erythrocytic, blood, and sexual stages. All recent approved malaria vaccines target the *P. falciparum* circumsporozoite protein (PfCSP) at the pre-erythrocytic stage to prevent human infection^2,3^. However, for malaria elimination, transmission-blocking vaccines are important tools to control parasite transmission from humans to mosquitos. Targeting the sexual stages of malaria parasites in the mosquito to block transmission is advantageous compared to other life cycle stages, as the development of the parasite in the mosquito entails bottlenecks in terms of parasite numbers^4^. Transmission-blocking immunogens and vaccines have been combined with those targeting pre-erythrocytic and blood stage parasites to generate multistage vaccines^5–10^. Such approaches have the potential to prevent both human infection and mosquito transmission, as well as limit the spread of escape mutants.

Two leading transmission-blocking vaccine candidates for *P. falciparum* are Pfs230 and Pfs48/45, and both are members of the 6-cysteine protein family. In *Plasmodium*, 6-cysteine proteins are some of the most abundant surface antigens and are expressed throughout the life cycle, with critical roles in parasite transmission, evasion of the host immune response and host cell invasion^11,12^. The defining feature of the family is the 6-cysteine domain^11,12^, which is conserved across apicomplexan parasites of the *Aconoidasida* class. This domain adopts a β-sandwich configuration, stabilized by up to six cysteines that form disulfide bonds^11,13,14^. With 14 6-cysteine domains (D1 – D14) and a molecular weight exceeding 300 kDa, Pfs230 is the largest member of the 6-cysteine family^14,15^ and was the first member of the family to be characterized, noteworthy for its capacity to elicit antibodies that block transmission to mosquitoes^16^. It is localized to the surface of gametocytes and gametes^17–19^ and has a critical role in male fertility, with male *Pfs230* gene knockouts unable to bind to red blood cells or form exflagellation centers in *P. falciparum*^20^ and unable to recognize female gametes in *P. berghei*^18^.

Pfs230 is thought to be localized to the parasite membrane via its interaction with Pfs48/45^21,22^, which has a putative glycosylphosphatidylinositol (GPI)-anchor^13,23^. Pfs48/45 has three 6-cysteine domains^14,15^ and shows a very similar localization pattern to Pfs230 on the surface of gametocytes and gametes^17–19,23–26^. When the *P48/45* gene is knocked out in *P. falciparum* and *P. berghei*, males are unable to attach to female gametes, leading to reduced ookinete production^18,26,27^. The crystal structure of the extracellular domain of Pfs48/45 has been determined^28^ and transmission blocking epitopes identified on all three 6-cysteine domains^29–33^. In addition, there are three ongoing Phase 1 clinical trials that use the Pfs48/45 antigen as a vaccine immunogen component^10,34,35^.

With respect to Pfs230, Pfs230D1-ExoProtein A (EPA) is the most clinically advanced transmission-blocking candidate^36^ and is currently undergoing a Phase 2 clinical trial in Mali (NCT03917654). Pfs230D1-EPA includes a part of the N-terminal pro-domain and domain 1 (D1) of Pfs230^37^. Full-length Pfs230 has never been produced recombinantly, and most transmission-blocking Pfs230 vaccine immunogens have focused on D1^10,37–45^. Crystal structures have been solved only for the D1 and D2 of Pfs230 ^42,43,46–48^. Therefore, the capability of the other 13 Pfs230 domains to function as transmission-blocking vaccine candidates is unknown. The expression of properly folded recombinant protein is essential for the characterization of these domains and the generation of specific antibodies to target them. However, the expression of correctly folded protein with sufficient yields for functional and structural studies has proved challenging.

Two key questions arise: how does Pfs230 interact with Pfs48/45, and are there additional domains that can function as vaccine candidates to block transmission? Here, by employing cryo-EM approaches, we present the structure of the endogenous Pfs230-Pfs48/45 fertilization complex and identify domains 13 and 14 of Pfs230 to be the critical interaction site for complex formation. We show how these domains are crucial for the localization of Pfs230 on activated gametes, nanobodies to these domains block complex formation and reduce oocyst numbers, and Pfs230 D13D14 mRNA-LNP vaccines induce specific antibodies against these domains. Overall, our study reveals novel transmission-blocking vaccine candidate domains against malaria parasite transmission.

### Cryo-EM structure of the Pfs230-Pfs48/45 fertilization complex shows the critical sites for complex formation

To investigate the structure of the Pfs230-Pfs48/45 fertilization complex, we purified the endogenous parasite proteins from late-stage sexual stage parasites and combined it with cryo-EM structure determination. We used a transgenic line Pfs230FL which has Pfs230 tagged with a C-terminal 3xFLAG-TwinStrepII (Figure S1). Using affinity pulldown followed by size exclusion chromatography (SEC), we successfully isolated the Pfs230-Pfs48/45 complex (Figure 1A and 1B). Pfs230 is present as 360 kDa and 310 kDa bands, with the lower band representing proteolytically cleaved Pfs230 (Figure 1B). Pfs48/45 migrates at ∼50 kDa as predicted from its molecular weight. Furthermore, we confirmed purification of the endogenous Pfs230-Pfs48/45 fertilization complex in multiple preparations using mass spectrometry (Figure S2).

**Figure 1.**
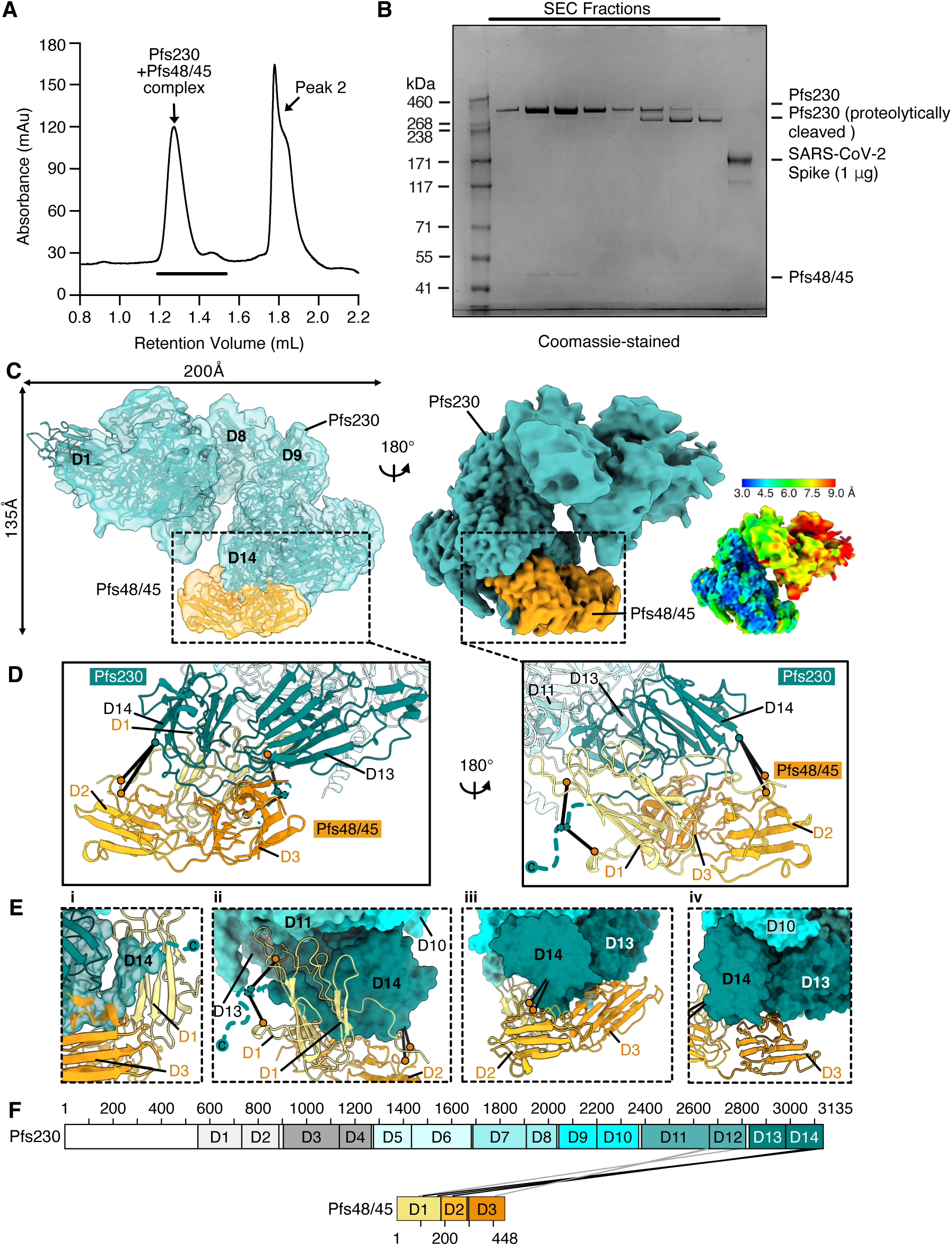
Cryo-EM structure of the Pfs230-Pfs48/45 fertilization complex shows critical sites for complex formation. (A) Size exclusion chromatography (SEC) profile of the purified Pfs230-Pfs48/45 complex from sexual stages of the malaria parasite. Peak 2 samples did not show any distinct protein bands on SDS-PAGE (data not shown). (B) SDS-PAGE analysis of purified Pfs230-Pfs48/45 complex corresponding to SEC fractions at retention volume 1.2-1.5 mL as indicated (panel A, black line). 1 μg of SARS-CoV-2 Spike protein was used to estimate the yield of Pfs230-Pfs48/45 complex. Pfs230 and Pfs48/45 are indicated, and molecular weight markers are on the left. (C) Composite map of the Pfs230-Pfs48/45 fertilization complex shown in two different views. The composite map was generated using the consensus map processed with EMReady (v2.0) and the local refinement map of the N-terminal subregion of Pfs230 (spanning D1-D8). Densities corresponding to Pfs230 (teal) and Pfs48/45 (orange) are highlighted. Left, cartoon representation of the structure placed into the cryo-EM map. Right, composite cryo-EM map colored by local resolution. (D) The Pfs230-Pfs48/45 binding site in two different orientations. Crosslinks consistent with the Pfs230-Pfs48/45 structure with a Cα-Cα distance less than 20 Å are indicated as black lines with orange and teal circles. Pfs230 residues that could not be placed with confidence into the density map are indicated with a dashed line in D and E. (E) Close-up views of the Pfs230-Pfs48/45 binding site with Pfs230 in surface representation and Pfs48/45 in cartoon representation. (F) Inter-molecular crosslinks between Pfs230 and Pfs48/45 after treatment with crosslinker sulfo-SDA. Crosslinks consistent with the Pfs230-Pfs48/45 structure with a Cα-Cα distance less than 20 Å are indicated by black lines and shown on the structures in D and E. Long distance crosslinks with a Cα-Cα distance larger than 20 Å are indicated by grey lines.

The cryo-EM map of the endogenous Pfs230-Pfs48/45 complex was determined to a global resolution of ∼3.36 Å (Figure 1C, Figure S3, Figure S4 and Table S1). The Pfs230-Pfs48/45 fertilization complex has a 1:1 stoichiometry and is approximately 200 Å wide and 135 Å long (Figure 1C). Pfs230 can be divided into two regions, an extended N-terminal region encompassing D1 to D8 which was solved with lower resolution in our EM map and a higher resolution C-terminal region consisting of D9 to D14, which is more compact and rigid as indicated by the local resolution estimation (Figure 1C, right).

Our cryo-EM structure reveals that Pfs48/45 docks into the bottom region of Pfs230 (Figure 1C). All three domains of Pfs48/45 interact with Pfs230, specifically engaging with its D13 and D14 domains, with a buried surface area of 2554 Å^2^ (Figure 1D). Notably, the C-terminal end of Pfs230 extends between the opening of Pfs48/45 D1 and D3 (Figure 1D and 1E, panel i). Among the three 6-cysteine domains of Pfs48/45, D1 is the most well-resolved with clear electron density for ∼98% of main chain residues. It binds to one side of Pfs230 D14, with three of its loops pointing towards Pfs230 D11 (Figure 1E, panel ii). Pfs48/45 D2 interacts with D14 (Figure 1E, panel iii), while D3 contacts both D13 and D14 (Figure 1E, panel iii and iv).

The interaction of Pfs230 with Pfs48/45 was further validated using crosslinking mass spectrometry, which showed that Pfs48/45 is in close proximity to the C-terminal end of Pfs230 (Figure 1F, Figure S5 and Data S1). Overall, 415 of 494 observed crosslinks were mapped on the structural model. Despite the high flexibility within the complex, about 82% of the crosslinks agree with the structural model with a median Cα-Cα distance of less than 20 Å (Data S1). There were four crosslinks found between Pfs230 and Pfs48/45, which are consistent with our Pfs230-Pfs48/45 cryo-EM structure. Intermolecular crosslinks with a Cα-Cα distance less than 20 Å were mapped onto the Pfs230-Pfs48/45 structure, showing that Pfs230 D14 is near Pfs48/45 D1 and D2 (indicated as black lines with orange and teal circles, Figure 1D and 1E, panels ii and iii). Collectively, these results clearly show that Pfs230 D13 and D14 are critical domains that interact with Pfs48/45.

### Tertiary fold of the endogenous Pfs230-Pfs48/45 complex

Our cryo-EM structure of Pfs230 is the first structure that contains multiple tandem pairs of 6-cysteine domains within one protein and provides an unprecedented opportunity to understand how these domains form their tertiary fold (Figure 2A). The 6-cysteine domain folds into a β-sandwich and has up to six cysteines that form disulfide bonds. For the *P. falciparum* 6-cysteine protein family, members have two, three or twelve 6-cysteine domains, with Pfs230 as the only member with 14 domains^11,12^. The 14 domains of Pfs230 form seven tandem pairs of 6-cysteine domains (D1D2, D3D4, D5D6, D7D8, D9D10, D11D12 and D13D14) (Figure 2B).

**Figure 2.**
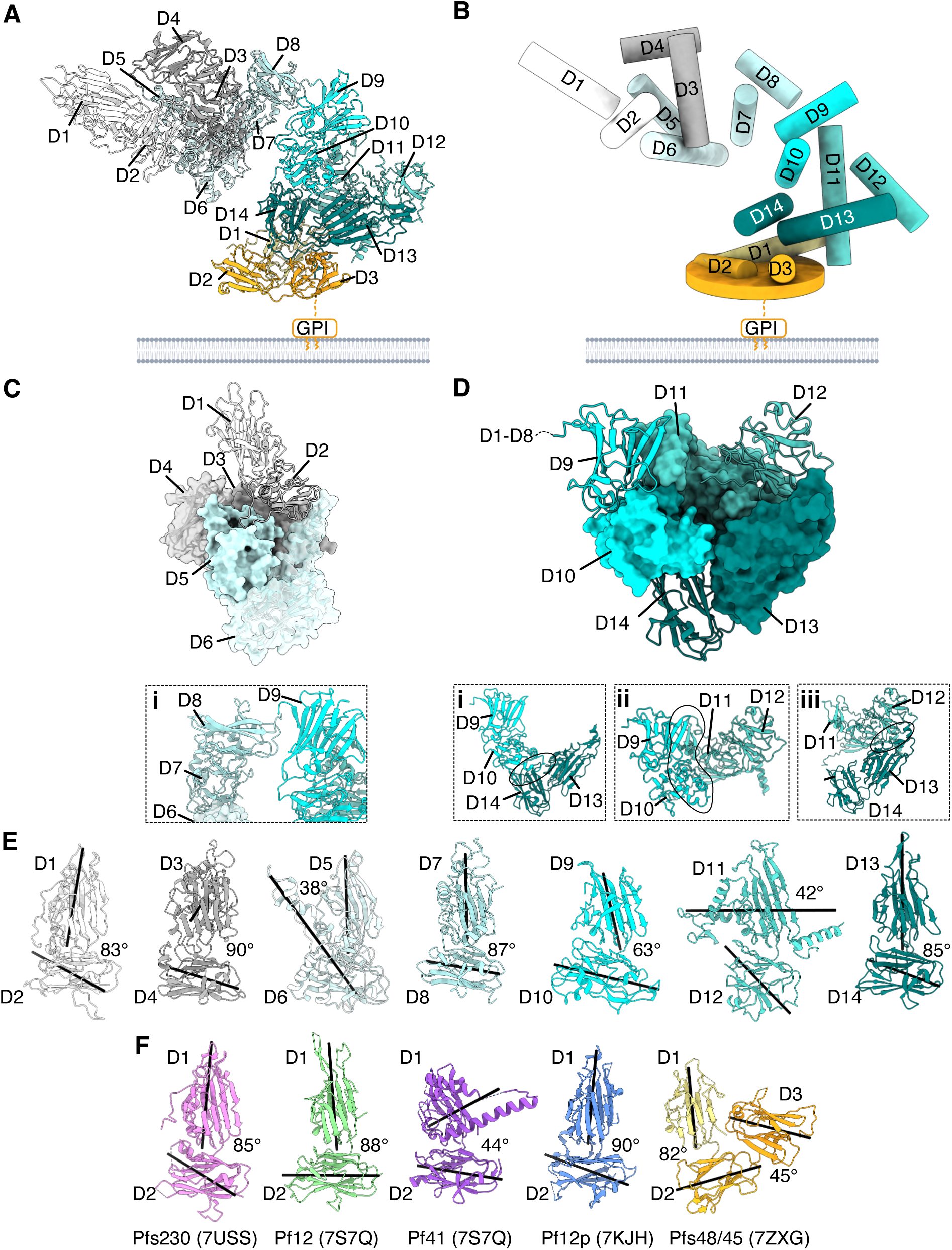
Tertiary fold of the endogenous Pfs230-Pfs48/45 fertilization complex. (A) Structure of the Pfs230-Pfs48/45 complex with the individual 6-cysteine domains indicated. Pfs230 and Pfs48/45 are colored in shades of teal and orange, respectively. The glycosylphosphatidylinositol anchor of Pfs48/45 is shown as GPI. (B) Simplified model of the Pfs230-Pfs48/45 complex organization. The center of mass of each 6-cysteine domain is represented by a centroid. (C) Stacking of multiple consecutive 6-cysteine domains in Pfs230 D1 to D6 with panel i showing a close-up view of Pfs230 D6 to D9. (D) Stacking of multiple consecutive 6-cysteine domains in Pfs230 D9 to D14 with panels i-iii showing the contacts of the central 6-cysteine domains D10, D11 and D13. (E) Tandem 6-cysteine domains of Pfs230 were aligned and shown in the same orientation next to each other. The center of mass of each domain is indicated by a black line and angles between tandem domain pairs are indicated. (F) Previously published crystal structures of 6-cysteine proteins containing two or three 6-cysteine domains with the PDB ID indicated. The center of mass of each domain and angles between tandem domain pairs are indicated.

At the globular level, the N-terminal (D1 - D8) and C-terminal regions (D9 - D14) of Pfs230 are arranged in a L-shape form to each other (Figure 2A-B). Within the N-terminal region of Pfs230, which encompasses D1 to D8, the tandem pairs D3D4 and D5D6 are arranged in a compact manner with the four domains forming a platform on which D2 sits (Figure 2C). Pfs230 D1 is positioned at the tip and is in contact only with D2. Furthermore, Pfs230 D7 is only in direct contact with its adjacent 6-cysteine domains D6 and D8, and D8 is only in direct contact with D7 and D9 (Figure 2C, panel i). For the C-terminal region, Pfs230 domains D9-D14 fold into a compact and rigid arrangement involving all three tandem pairs, D9D10, D11D12 and D13D14 (Figure 2D). Pfs230 D10, D11 and D13 form the ‘center’ and are surrounded by four other 6-cysteine domains each: Pfs230 D10 is surrounded by D9, D11, D13 and D14; Pfs230 D11 is surrounded by D9, D10, D12 and D13; and Pfs230 D13 is surrounded by D10, D11, D12 and D14 (Figure 2D). Notably, each of the three domains in the center (D10, D11, D13) contacts both domains of a neighboring tandem pair: D10 contacts both D13 and D14; D11 contacts both D9 and D10; and D13 contacts both D11 and D12 (Figure 2D, panel i-iii, respectively).

The two β-sandwich domains of each tandem pair are connected by a short linker and tilted against each other with varying degrees from 38° to 90° based on their center of mass (Figure 2E). We observe a similar range of interdomain tilting from crystal structures of other 6-cysteine proteins that contain tandem pairs such as Pf12, Pf41, Pf12p and Pfs48/45 (Figure 2F)^46,49–53^.

### Domains 13 and 14 of Pfs230 are critical for the localization of Pfs230 on *P. falciparum* gametes and successful parasite fertilisation

Pfs230 is found on the surface of gametes but lacks a membrane anchor. For its surface localization, it is proposed that Pfs230 forms a stable complex with Pfs48/45, which contains a putative GPI anchor^13,20–23^. Our cryo-EM shows that D13 and D14 of Pfs230 are required for the interaction with Pfs48/45, therefore we wanted to determine if the removal of those domains would perturb Pfs230 localization on gametes. We generated a *P. falciparum* transgenic line that expresses Pfs230 without D13 and D14 (230Δ1314, Figure S6A) in a NF54/iGP2 background^54^. As with Pfs230FL, 230Δ1314 was also tagged with a C-terminal 3xFLAG-TwinStrepII (Figure S6A). Using an anti-Pfs230 monoclonal antibody (mAb) LMIV230-01^42^, we detected a 350 kDa band for 230FL stage V gametocytes under non-reducing conditions (Figure 3A). In contrast, Pfs230 migrated as a lower molecular weight band in the 230Δ1314 line, which is expected due to the removal of amino acids spanning K2828 to L3135, resulting in a loss of 36 kDa. The respective bands in Pfs230FL and 230Δ1314 were also detected using anti-FLAG and anti-StrepII antibodies (Figure 3A). These results show that Pfs230 without D13 and D14 is still expressed in late-stage gametocytes.

**Figure 3.**
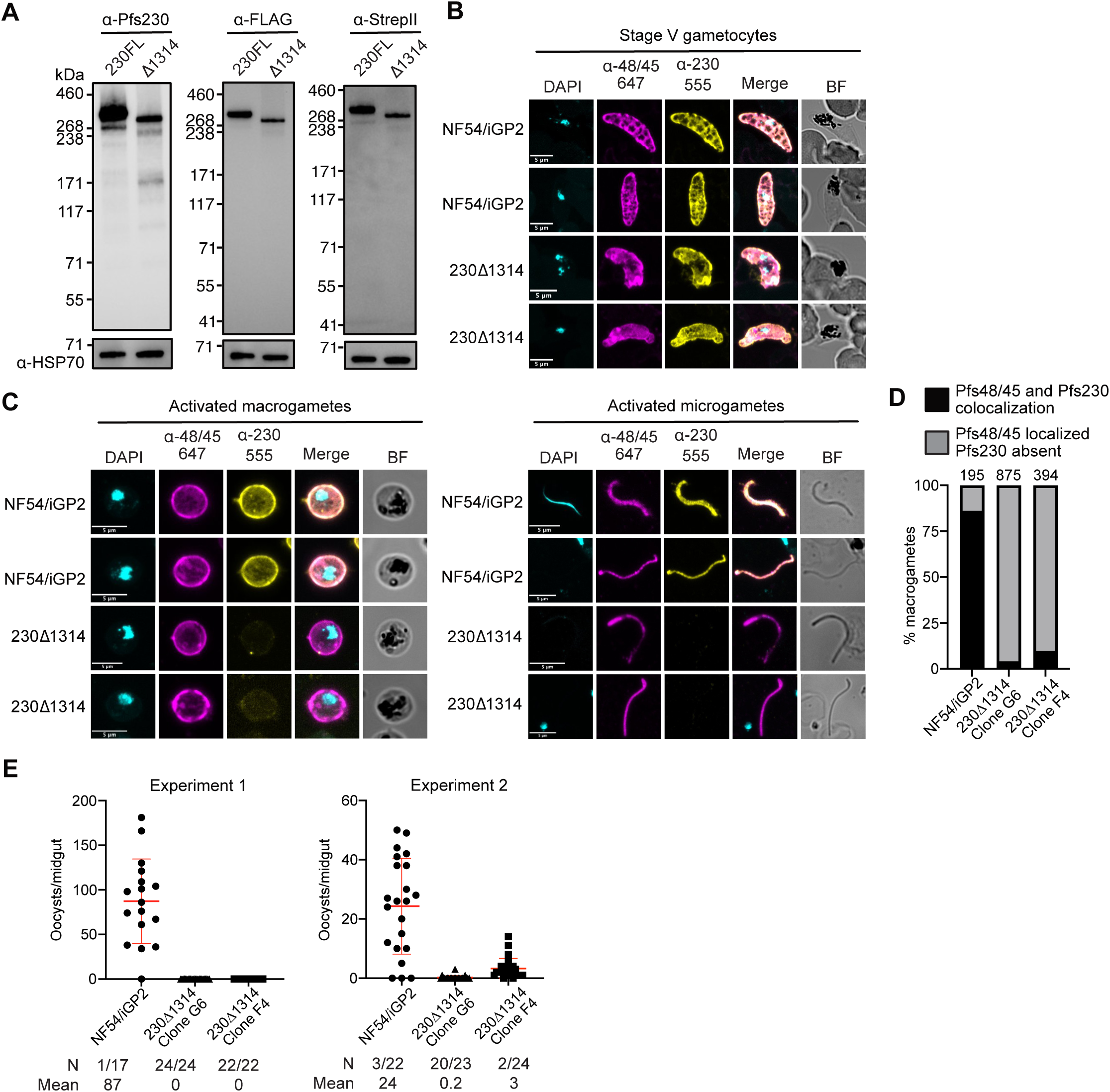
Domains 13 and 14 of Pfs230 are critical for the localization of Pfs230 on *P. falciparum* gametes. (A) Western blotting analyses using anti-Pfs230 LMIV230-01, anti-FLAG and anti-StrepII antibodies show that 230ΔD13D14 line expresses a truncated Pfs230, which runs at a lower molecular weight than the full-length protein present in 230FL. Both lines express C-terminal 3xFLAG and TwinStrepII tags. Anti-HSP70 was used as a loading control. All samples were run in non-reducing conditions. Molecular weight markers are shown on the left. (B) Immunofluorescence assay of stage V gametocytes from NF54/iGP2 expressing full-length Pfs230 or 230Δ1314. Fixed gametocytes were stained with anti-Pfs48/45 TB31F conjugated to Alexa 647 and anti-Pfs230 LMIV230-01 directly conjugated to Alexa 555. Parasite nuclei are stained with DAPI (cyan), merged and bright field (BF) images are shown. Scale bar = 5 μm. (C) Surface immunofluorescence assay (SIFA) of activated macrogametes (left panel) and microgametes (right panel) of NF54/iGP2 which express full length Pfs230 and two clonal lines of 230ΔD13D14. Gametes were stained with anti-Pfs48/45 TB31F conjugated to Alexa 647 and anti-Pfs230 LMIV230-01 directly conjugated to Alexa 555 prior to fixation. Merged and bright field (BF) images are shown. Scale bar = 5 μm. (D) Automated quantitation of Pfs230 localization observed in the SIFA of activated macrogametes, where anti-Pfs230 staining was classified as either being colocalized with anti-P48/45 staining (black) or absent while Pfs48/45 was staining was present (grey). The number of activated macrogametes counted for each parasite line is shown above each stacked bar graph. (E) SMFA assessing the transmissibility of NF54/iGP2 and two clonal lines of 230Δ1314 (G6 and F4). Number of oocysts per dissected midgut is plotted for two experiments. Bars show the mean and standard deviation. N, number of uninfected mosquitoes / total number of mosquitoes.

Malaria parasite fertilization occurs in the female *Anopheles* mosquito midgut and Pfs230 and Pfs48/45 are critical for successful fertilization^18,20,26,27^. Mature male and female gametocytes are activated in the mosquito midgut by environmental changes such as temperature drop, increased pH, and the presence of mosquito xanthurenic acid^55–58^. Upon activation, gametes egress from red blood cells for parasite fertilization and subsequent zygote formation. We initially compared the localization of Pfs230 and Pfs48/45 in stage V gametocytes between the parental NF54/iGP2 and 230Δ1314 lines. Using an immunofluorescence assay with anti-Pfs48/45 mAb TB31F^53^ and anti-Pfs230 mAb LMIV230-01^42^, which were directly conjugated to Alexa647 and Alexa555, respectively, we observed colocalization of Pfs230 and Pfs48/45 in stage V gametocytes in both NF54/iGP2 and 230Δ1314 line (Figure 3B). However, we observed that the lattice-like pattern of Pfs230 and Pfs48/45 staining seen in the NF54/iGP2 stage V gametocytes was absent in the 230Δ1314 line (Figure 3B). These immunofluorescence results show that Pfs230 is still expressed in 230Δ1314 line and colocalizes with Pfs48/45. However, since stage V gametocytes are still within the red blood cells, we are unable to determine if Pfs230 is surface localized without D13 and D14.

Next, to determine if Pfs230 was localized on the surface of activated male and female gametes, we used a surface immunofluorescence assay (SIFA) (Figure 3C and Figure S6B). As expected for NF54/iGP2, Pfs230 and Pfs48/45 were expressed and colocalized on both male microgametes and female macrogametes (Figure 3C). In contrast, for 230Δ1314, while we observed Pfs48/45 localization on the surface of microgametes and macrogametes, there was an absence of Pfs230 on the surface of the gametes (Figure 3C). In two independent clones of 230Δ1314, we show that over 90% of activated female gametes have no Pfs230 localization on their surfaces compared to 14% in NF54/iGP2 (Figure 3D and Figure S6C). These results clearly show that D13 and D14 of Pfs230 are critical for its localization on the surface of male microgametes and female macrogametes. We also examined if the 230Δ1314 transgenic parasites had a defect in parasite fertilization. Using the standard membrane feeding assay (SMFA), we observed that NF54/iGP2 had a mean oocyst count of 87 and 24 oocysts per midgut in experiments 1 and 2 respectively (Figure 3E). However, in two independent clones of 230Δ1314, we observed a significant reduction of oocyst formation with a mean oocyst count of 0 per midgut in experiment 1, and mean oocyst count of 0.2 and 3 oocysts per midgut in experiment 2 (Figure 3E). These results clearly show that D13 and D14 of Pfs230 are critical for its role in parasite fertilization.

### Nanobodies specific to Pfs230 D13D14 recognize gametocytes

We wanted to examine if nanobodies against Pfs230 D13D14 could inhibit Pfs230-Pfs48/45 complex formation. Using phage display, we identified 11 unique nanobodies (W2802 – W2812) using Sanger sequencing and based on differences in the amino acid sequence of the complementary determining region 3 (CDR3) that vary by at least one amino acid (Figure 4A and Data S2). All these unique antibodies were also identified among the top 100 most abundant sequences by Next-Generation Sequencing (NGS) after two rounds of phage display, with consistent enrichment throughout the panning rounds (Figure 4B and Figure S7). Seven of these nanobodies (W2806 – W2812) bound to Pfs230 D13D14 by ELISA (Figure 4A). In addition, nanobodies F5 and F10 raised against Pfs230 D1D2^46^ and nanobodies targeting other 6-cysteine proteins, such as anti-Pf12p nanobody C12 and anti-Pf41 nanobody G12, had no detectable binding to Pfs230 D13D14^51,52^ (Figure 4A).

**Figure 4.**
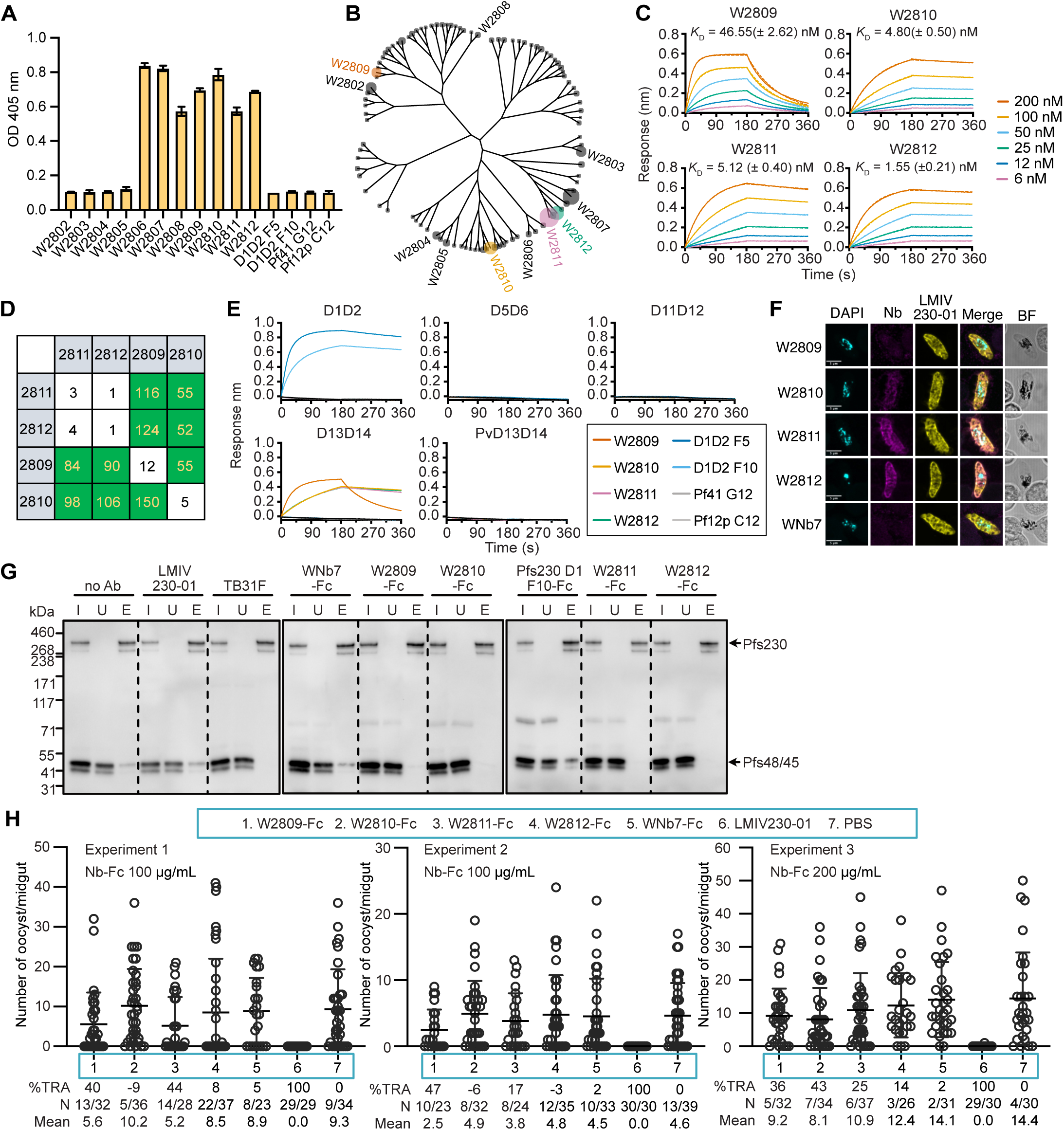
Nanobodies specific to Pfs230 D13D14 recognize gametocytes, block complex formation and reduce oocyst numbers in mosquitoes. (A) Nanobody binding to Pfs230 D13D14 was evaluated by ELISA. Error bars represent standard deviation of the mean of two technical replicates. (B) Cladogram of the 100 most abundant nanobodies from Next-Generation Sequencing (NGS) after two rounds of phage display selection. Pfs230 D13D14 nanobodies identified by Sanger sequencing are labelled. Tree tips are scaled relative to abundance (counts per million). (C) Biolayer interferometry (BLI) affinity measurements using a two-fold dilution series of recombinant Pfs230 D13D14 from 6 – 200 nM binding to immobilized nanobodies. Measurements were plotted (solid line) and fitted to a 1:1 binding model (dashed line). Mean *K*_D_ values for replicates with standard deviations are indicated, and representative binding curves are shown from two independent experiments. (D) Epitope binning competition experiments using BLI with the left column indicating immobilized nanobodies and the top row indicating nanobodies pre-incubated with Pfs230 D13D14 at a 10:1 molar ratio. Binding of Pfs230 D13D14 premixed with nanobody was calculated relative to Pfs230 D13D14 binding alone, which was arbitrarily assigned as 100%. The green and white boxes represent non-competition and competition, respectively. (E) Binding specificity of nanobodies to different Pfs230 domains and Pvs230 D13D14 by BLI. Representative curves of two independent experiments. (F) Immunofluorescence assay of stage V gametocytes using the indicated nanobody (Nb) detected with goat anti-alpaca IgG VHH domain conjugated to Alexa Fluor 488 (magenta) and anti-Pfs230 LMIV230-01 antibody conjugated to Alexa 555 (yellow). Parasite nuclei are stained with DAPI (cyan), merged and brightfield (BF) images are shown. Scale bar = 5 μm. (G) Pull down of Pfs230-Pfs48/45 complex from parasite lysate using StrepTactin XT resin in the presence of various antibodies or nanobody-Fcs. Presence of the Pfs230-Pfs48/45 complex was examined using western blotting under non-reducing conditions. Pfs48/45 and Pfs230 are indicated on the right. Representative blots of n = 3 independent experiments. I, input; U, unbound; E, eluate. Molecular weight ladder labeled on the left in kDa. (H) Transmission-blocking activity of anti-Pfs230 D13D14 nanobody-Fcs by standard membrane feeding assay. Number of oocysts per dissected midgut are plotted for three experiments. For experiments 1 and 2, nanobody-Fcs were added to blood meals at a concentration of 100 µg/mL. For experiment 3, nanobody-Fcs were added at a concentration of 200 µg/mL. LMIV230-01 was added at 200 µg/mL in all experiments. Bars show means with standard deviation. TRA, transmission reducing activity; N, number of uninfected mosquitoes / total number of mosquitoes.

Of the seven Pfs230 D13D14 nanobodies, four nanobodies (W2809, W2810, W2811 and W2812) bound to recombinant Pfs230 D13D14 as determined by biolayer interferometry (BLI), with affinities from 1.6 to 46.6 nM (Figure 4C and Data S2). W2811 and W2812 bound Pfs230 in the same epitope bin, while W2809 and W2810 bound distinct epitopes to the others (Figure 4D). To examine the specificity of the four Pfs230 D13D14 nanobodies, we expressed and purified Pfs230 D1D2, D5D6 and D11D12, which have 20%, 18% and 22% amino acid sequence identity to Pfs230 D13D14, respectively. We also expressed the homologous D13D14 domains in Pvs230. The four anti-Pfs230 D13D14 nanobodies did not recognize Pfs230 domains D1D2, D5D6 or D11D12 or Pvs230 D13D14 by BLI (Figure 4E). Collectively, these results show that W2809, W2810, W2811 and W2812 are highly specific for Pfs230 D13D14. Pfs230 nanobodies W2810, W2811 and W2812 staining colocalize with anti-Pfs230 mAb LMIV230-01 staining on fixed stage V gametocytes (Figure 4F). However, there was no detectable localization of Pfs230 in gametocytes using nanobody W2809, which may suggest its binding is sensitive to the fixation methods used. As expected, SARS-CoV-2 nanobody WNb7^59^ did not show any detectable staining of gametocytes (Figure 4F).

### Structural analyses of Pfs230 D13D14 nanobodies that inhibit Pfs230-Pfs48/45 complex formation and have transmission-reducing activity

To determine if anti-Pfs230 nanobodies inhibit complex formation, we performed pull-down assays to isolate the endogenous Pfs230-Pfs48/45 complex using our transgenic 230FL line. Endogenous Pfs230-Pfs48/45 complex was pulled down using StrepTactin XT resin in the absence of any antibody, as observed in the eluate lane (lane E, no Ab) (Figure 4G). In the presence of anti-Pfs230 mAb LMIV230-01, anti-SARS-CoV-2 nanobody-Fc fusion WNb7-Fc and anti-Pfs230 D1D2 nanobody-Fc fusion F10-Fc, we still observed the presence of the Pfs230-Pfs48/45 complex in the eluate lane, showing that these antibodies do not inhibit complex formation. In contrast, the addition of anti-Pfs48/45 mAb TB31F resulted in the loss of Pfs48/45, indicating that complex formation between Pfs230 and Pfs48/45 was disrupted (Figure 4G). We observed a similar loss of Pfs48/45 with the addition of the anti-Pfs230 D13D14 nanobody-Fc fusions W2809-Fc, W2810-Fc, W2811-Fc and W2812-Fc, showing that these nanobodies can inhibit complex formation between Pfs230 and Pfs48/45 (Figure 4G).

To determine whether the ability of the nanobodies to block complex formation correlated with an ability to block parasite transmission, their transmission-blocking capability was assessed by SMFA. LMIV230-01 was used as a positive control and consistently showed >99% transmission-reducing activity (TRA) at a concentration of 200 µg/mL (Figure 4H), as previously reported^42,46^. PBS and WNb7-Fc were used as negative controls. At 100 µg/mL, nanobody-Fc fusion W2809-Fc reduced oocyst numbers in the mosquito across two repeat experiments, showing TRA of up to 47% (Figure 4G, left and middle panels). However, increasing the concentration of nanobody-Fc to 200 µg/mL did not increase TRA for W2809-Fc (Figure 4H, right panel). W2811-Fc showed TRA of between 17% and 44% across experiments; W2810-Fc did not show any TRA at 100 µg/mL but showed TRA of 43% at 200 µg/mL; and W2812-Fc did not show any TRA across experiments (Figure 4H).

To understand the interaction of these nanobodies with Pfs230, we determined the crystal structures of the Pfs230 D13D14-W2809 complex, Pfs230 D13D14-W2810 complex and Pfs230 D13D14-W2812 complex at resolutions of 2.5 Å, 1.9 Å, and 3.2 Å, respectively (Figure 5A and Table S2). The crystal structures show that nanobodies W2810 and W2812 bind to D13 exclusively, whereas W2809 binds both domains, with most contacts formed with D14 (Figure 5A, Figure 5B and Table S3). All three nanobodies bind to non-overlapping regions, as predicted by the epitope binning experiments (Figure 4D). W2809, W2810 and W2812 bind to Pfs230 D13D14 with a buried surface area of 711 Å^2^, 588 Å^2^ and 685 Å^2^, respectively (Figure 5B). The W2809 epitope overlaps where Pfs48/45 D1 predominantly engages with Pfs230 D14 (Figure 5C). While the epitopes of W2810 and W2812 do not overlap with the Pfs230-Pfs48/45 interaction site, their binding may disrupt the conformation or interactions of D13 with neighboring Pfs230 domains, such as D12 and parts of D11 (Figure 5C). Overlaying our cryo-EM structure of Pfs230-Pfs48/45 with the crystal structure of TB31F Fab bound to Pfs48/45 (PDB ID 6E63)^53^ shows that the TB31F epitope overlaps with where Pfs230 D13D14 binds Pfs48/45 D3 (Figure 5D and E).

**Figure 5.**
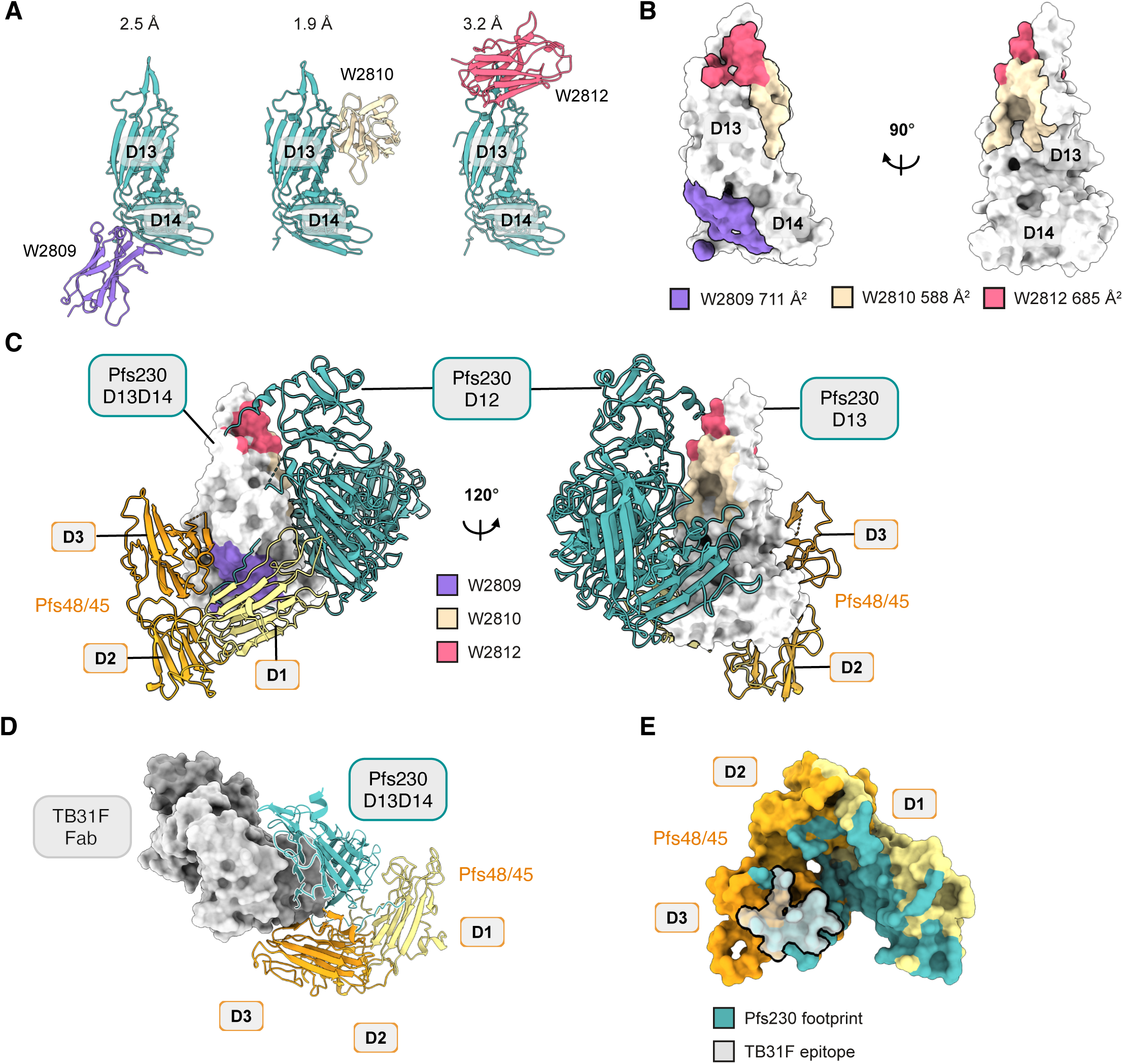
Structural analyses of Pfs230 D13D14 nanobodies that inhibit Pfs230-Pfs48/45 complex formation. (A) Pfs230 D13D14/W2809 complex at 2.5 Å, Pfs230 D13D14/W2810 complex at 1.9 Å and Pfs230 D13D14/W2812 complex at 3.2 Å resolution. Pfs230 D13D14 is shown in teal and nanobodies W2809, W2810 and W2812 are shown in purple, beige and pink, respectively. (B) Surface representation of Pfs230 D13D14 showing the epitope footprint of residues within 4 Å of the nanobodies, with W2809, W2810 and W2812 shown in purple, beige and pink, respectively. (C) Cryo-EM structure of Pfs230 D9 to D14 and Pfs48/45 is shown in two orientations, with Pfs230 domains D13 and D14 shown in surface representation (white) and domains D9 to D12 in cartoon representation (teal). The epitope footprint of W2809, W2810 and W2812 is shown in purple, beige and pink, respectively. (D) TB31F and Pfs230 clash for binding to Pfs48/45. Overlay of PDB ID 6E63 with cryo-EM structure of Pfs230 D13D14-Pfs48/45. TB31F Fab is shown in grey in surface representation. (E) Overlapping binding sites of Pfs230 and TB31F Fab on D3 of Pfs48/45. Pfs48/45 is shown in surface representation with Pfs230 and TB31F footprints in teal and light grey, respectively.

### Pfs230 D13D14 is a target of naturally acquired antibodies

To determine if Pfs230 domains are targets of naturally acquired antibodies, we used a multiplexed assay^60^ to screen for reactivity of naturally acquired humoral immunity to Pfs230 domains and its ortholog from *P. vivax*, Pvs230, in children from Papua New Guinea (PNG) where *P. falciparum* and *P. vivax* are hyperendemic. We found that the mean total IgG antibodies against Pfs230 D1D2, D5D6, D11D12 and D13D14 were significantly higher in PNG children than in malaria-naïve individuals (Figure 6A and 6B). Our finding suggests that individuals living in malaria-endemic areas can develop immune responses to Pfs230 and Pvs230 through exposure to natural infections. Interestingly, the mean IgG levels against most constructs were notably higher in individuals with a current *P. falciparum* infection (Pf pos) as compared to those who had a prior infection (Pf prior), especially for Pfs230 D1D2 (Figure 6B). These findings suggest that antibodies induced by natural infection may not confer long-lived immunity in the absence of recurrent infections. This study proposes that a Pfs230 vaccine could potentially be employed as a booster to maintain high levels of Pfs230-specific antibody, especially in areas with low malaria transmission and/or that natural infections may effectively boost vaccine-induced responses against both D1D2 and D13D14 domains. Additionally, the immunoreactivity against Pv230 D13D14 in these samples with *P. falciparum* infections (Figure 6B) is indicative of potential cross-reactivity between the two species, as only 13 individuals had a concurrent *P. vivax* infection.

**Figure 6.**
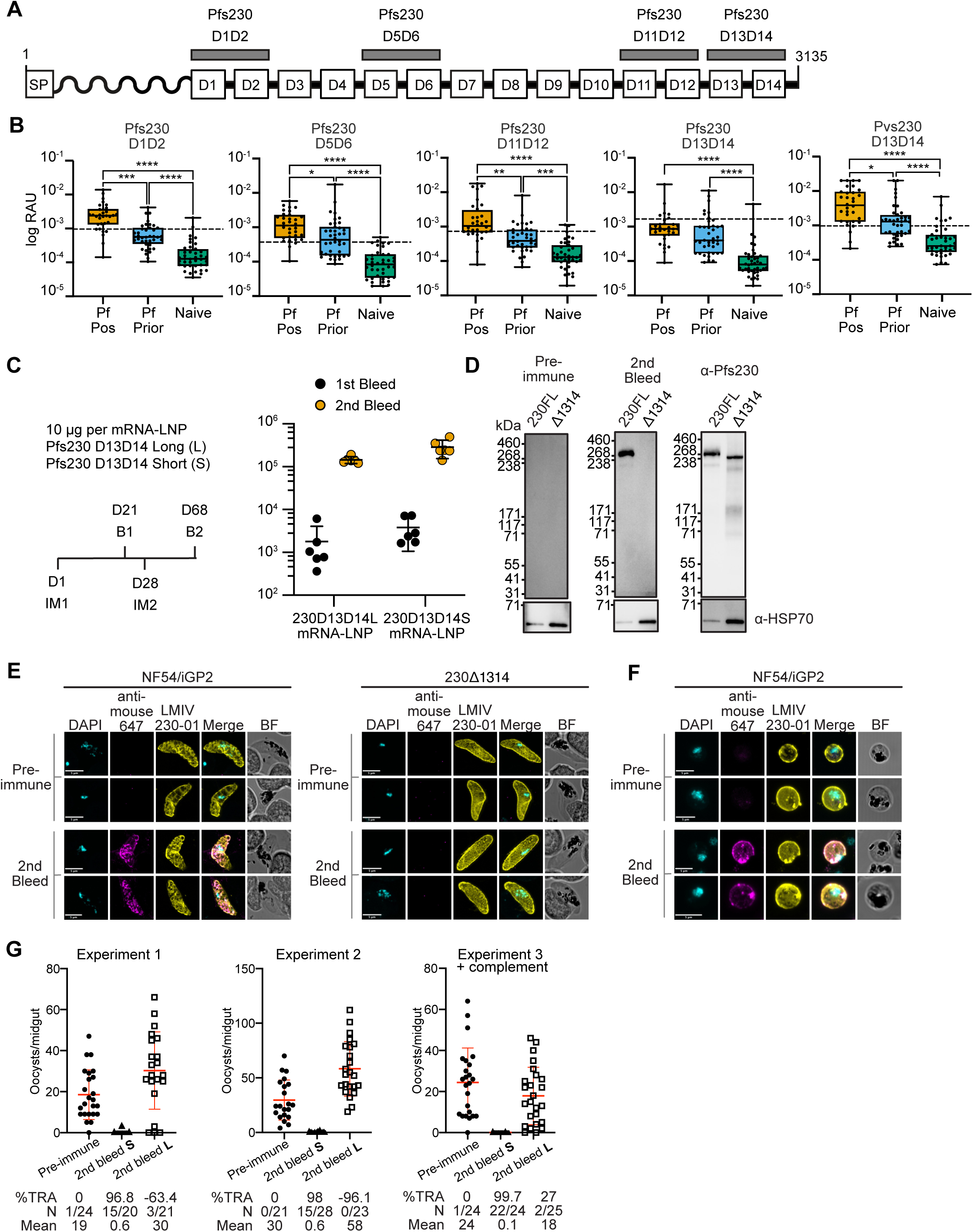
Pfs230 D13D14 is a target of naturally acquired immunity and a potential transmission-blocking vaccine candidate. (A) Schematic of full-length Pfs230 with the recombinant fragments to D1D2, D5D6, D11D12 and D13D14 indicated. Signal peptide is labeled as SP. (B) Multiplex assay for reactivity of plasma against Pfs230 and Pvs230 domains from PNG individuals with current *P. falciparum* infection (Pf pos) as compared to those who had a prior infection (Pf prior) or naïve individuals from Melbourne. The dashed line represents an antigen-specific seropositivity cut-off, calculated as the mean of the negative control samples plus two times the standard deviation. Statistical significance between groups was calculated using a Kruskal-Wallis test, with Dunn’s multiple comparisons test. * p ≤ 0.05, ** p ≤ 0.01, *** p ≤ 0.001, **** p ≤ 0.0001. (C) Groups of C57BL/6 mice were immunized with 10 μg of either 230D13D14L or 230D13D14S mRNA-LNP alone. Mice were immunized (IM) on day 1 and day 28. For pre-immune sera, mice were bled on day 1 prior to immunization. Mice were then bled on day 21 and day 68 for the first (B1) and second (B2) bleed. Pfs230 D13D14 antibody levels were measured by endpoint dilution ELISA. Error bars represent 95% confidence intervals of the geometric mean. n = 6 mice per immunization group. (D) Western blot analyses showing immunized mouse sera are specific to Pfs230 D13D14 in sexual stage parasites. Anti-HSP70 was used as a loading control. Molecular weight markers are shown on the left. (E) Immunofluorescence assay of stage V gametocytes from NF54/iGP2 expressing full-length Pfs230 and 230Δ1314. Fixed gametocytes were stained with pre-immune or second bleed pooled sera from mice immunized with 230D13D14S mRNA-LNP. (F) Surface immunofluorescence assay (SIFA) of activated macrogametes from NF54/iGP2. Macrogametes were stained with pre-immune or second bleed pooled sera from mice immunized with 230D13D14S mRNA-LNP. For panels E and F, mouse sera were detected using donkey anti-mouse conjugated to Alexa 647 and anti-Pfs230 LMIV230-01 was directly conjugated to Alexa 555. Parasite nuclei are stained with DAPI (cyan), merged and bright field (BF) images are shown. Scale bar = 5 μm. (G) SMFA using either pre-immune or second bleed following immunization with 230D1314S and 230D13D14L mRNA-LNP. Number of oocysts per dissected midgut are plotted for three experiments, experiment 3 included complement active human serum. Bars show means and standard deviation. TRA, transmission reducing activity; N, number of uninfected mosquitoes / total number of mosquitoes. Mean, mean number of oocysts per midgut.

### Pfs230 D13D14 mRNA-LNP elicit high antibody titers that block transmission

Two mRNA constructs for Pfs230 D13D14 were formulated as lipid nanoparticles (mRNA-LNP) (Figure 6C). The first mRNA encompassed a longer fragment from K2828-L3135 (230D13D14L) and the second encompassed a shorter fragment of Pfs230 D13D14 from K2828-E3111 (230D13D14S). Both constructs had a N-terminal secreted alkaline phosphatase signal peptide (SEAP) but no added transmembrane domain or GPI anchor. 230D13D14L and 230D13D14S mRNA-LNPs were used to immunize BALB/c mice and immune sera were assayed for antibody responses against recombinant Pfs230 D13D14 (Figure 6C and Figure S8). After the first immunization, 230D13D14L and 230D13D14S elicited end point antibody titers of 1787 and 3824, respectively. Administration of the second immunization of both constructs resulted in significant boosting of antibody titers 80-fold and 74-fold, with 230D13D14L and 230D13D14S eliciting end point antibody titers of 144263 and 284235, respectively (Figure 6C). As expected, we did not observe any immune response to Pfs230 D13D14 in the pre-immune sera from both sets of mice, showing that the mRNA-LNP immunizations were directly responsible for the generation of Pfs230-specific antibodies (Figure S8).

Due to the limited volume of sera collected, sera from the second bleed of each mouse were pooled to determine specificity of the antisera to parasite material. Using pooled sera from the 230D13D14S mRNA-LNP immunizations, we detected a 360 kDa band in 230FL stage V gametocytes under non-reducing conditions (Figure 6D, middle panel). In contrast, we did not detect any band in the 230Δ1314 line, showing that the antibodies elicited by 230D13D14S mRNA-LNP immunization are completely specific to D13D14 of Pfs230. As a control, mAb LMIV230-01 which binds to Pfs230 D1 could detect the 360 kDa band in 230FL stage V gametocytes and a lower molecular weight band in the 230Δ1314 line under non-reducing conditions (Figure 6D, right panel). Pre-immune mouse sera did not detect any band in either 230FL or the 230Δ1314 line (Figure 6D, left panel) showing that the antibodies against Pfs230 D13D14 were elicited via mRNA-LNP immunization.

The pooled sera from the 230D13D14S mRNA-LNP immunization were able to detect Pfs230 on both stage V gametocytes using an immunofluorescence assay (Figure 6E, bottom left panels) and the surface of macrogametes using SIFA (Figure 6F, bottom panels). However, the pooled immune sera did not detect Pfs230 in the 230Δ1314 clonal lines (Figure 6E, bottom right panels), again demonstrating the specificity of antibodies against D13 and D14 elicited by immunization. The anti-Pfs230 D13D14 sera staining pattern colocalized with that of the anti-Pfs230 mAb LMIV230-01 in wild-type NF54/iGP2 (Figure 6E and 6F). As expected, the pooled pre-immune sera did not show any detectable staining of stage V gametocytes or macrogametes (Figure 6E and 6F, top panels).

SMFAs were performed using the sera pooled from pre-immune mice and the 230D13D14S and 230D13D14L mRNA-LNP immunizations to determine whether antibodies generated could reduce transmission. In two experiments, we observed anti-Pfs230 D13D14S sera reduced transmission by 97 and 98% when compared to pre-immune sera (Figure 6G, experiment 1 and 2), whereas anti-Pfs230 D13D14L showed no reduction in transmission activity. In a third experiment, we included complement active human serum to assess how complement would affect transmission blocking activity. Here we observed similar results to the SMFA performed without active human complement. The pre-immune serum had a mean of 24 oocysts per midgut and the addition of anti-230D13D14S sera in the presence of active human complement reduced transmission by 99.7% (Figure 6G, experiment 3). Collectively, these results show that 230D13D14S mRNA-LNP immunization can elicit Pfs230 D13D14-specific antibodies that significantly reduce parasite transmission independent of active human complement.

## Discussion

The cryo-EM structure of the Pfs230-Pfs48/45 fertilization complex identified Pfs230 D13D14 as important for Pfs230’s interaction with Pfs48/45. Using a transgenic *P. falciparum* line that expresses Pfs230 without D13 and D14, we show that the presence of these domains is critical for the localization of Pfs230 on the surface of male microgametes and female macrogametes, and for parasite fertilization. Nanobodies against Pfs230 D13 and D14 inhibited Pfs230-Pfs48/45 complex formation, and several can reduce oocyst formation in the mosquito midgut. Crystal structures of the inhibitory nanobodies bound to Pfs230 D13D14 provide mechanistic insights on future approaches to develop more potent transmission-blocking antibodies. Pfs230 D13D14 is a target of naturally acquired immunity in PNG children with current and prior *P. falciparum* infection. In addition, we show that 230D13D14S mRNA-LNP immunogen can elicit Pfs230 D13D14-specific antibodies capable of reducing transmission in mosquitoes. This study proposes that a Pfs230 D13D14 vaccine could potentially be employed as a booster to maintain high levels of Pfs230 D13D14-specific antibodies, especially in areas with low malaria transmission. Collectively, our results show that Pfs230 D13D14 is a novel transmission-blocking vaccine candidate. Furthermore, Pfs230 D14 alone shows promise as a target of transmission-blocking antibodies or to be developed as a smaller subunit vaccine. Our cryo-EM structure shows that Pfs230 D14 is engaged with all three Pfs48/45 domains and its C-terminal end is sandwiched between D1 and D3 of Pfs48/45, supported by our cross-linking studies. In addition, nanobody W2809 which binds mostly to Pfs230 D14, blocks Pfs230-Pfs48/45 complex formation and reduces transmission by up to 47%. In the last 30 years, most Pfs230 vaccine immunogens have focused on the development of D1, in part driven by its ability to generate potent transmission-blocking antibodies and the ease of recombinant protein expression for this domain^10,37–45^. To date, it is still not understood how Pfs230 D1 functions in fertilization. While there have been previous studies expressing multiple domains of Pfs230, they have been hindered by low level expression, which precludes additional functional analyses beyond immunological studies^38,39,61–63^. More recently, it was discovered that a mouse monoclonal antibody 18F25.1 targets Pfs230 D7 and potently reduces *P. falciparum* transmission in a complement-dependent manner^64^, clearly showing that other Pfs230 domains can be novel transmission-blocking vaccine candidates. As such, our comprehensive structural analyses of the Pfs230-Pfs48/45 fertilization complex, the full-length endogenous Pfs230 and nanobody complexes facilitate future rational design of vaccines or monoclonal antibody approaches.

Our study shows that structural approaches with endogenous parasite material is a better suited approach to study the 6-cysteine protein family for which recombinant protein expression remains challenging across multiple expression systems. Despite the critical role of the 6-cysteine family in important functions such as parasite transmission, evasion of the host immune response and host cell invasion^11,12^, structures of the complete ectodomains of most 6-cysteine proteins are lacking, including major vaccine candidates Pfs47, PfLISP2, Pf36 and Pf52. Indeed, of the 14 members of the family in *P. falciparum*, structural information is only available for five, namely Pf12^50,52^, Pf12p^51^, Pf41^49,52^, Pfs48/45^28,53,65^ and Pfs230^42,43,46^. Cryo-EM studies like ours would also be beneficial for heterodimeric 6-cysteine proteins that bind human receptors. For example, Pf36 and Pf52 form a heterodimeric complex^66^ and Pf36 binds several different receptors such as host receptors EphA2^67^, CD81 and SR-B1^68^ to facilitate sporozoite invasion. Both of these proteins feature in a genetically attenuated parasite (GAP) vaccine, which has undergone a Phase I clinical trial (NCT03168854), and was capable of eliciting antibodies that inhibited sporozoite invasion and traversal^69^. Cryo-EM approaches may provide structural insights for understanding the interactions and molecular function of the 6-cysteine proteins and how they can be inhibited.

To the best of our knowledge, our nanobodies are the first collection of monoclonal antibodies to recognize Pfs230 D13D14. Over 20 inhibitory mAbs targeting *P. falciparum* Pfs230 have been reported, with transmission-reducing activities ranging from 42% to 100%^16,17,19,40,42,43,46,63,64,70,71^. All these mAbs bind to Pfs230 D1, except for 18F25.1, which binds to Pfs230 D7. Our nanobody W2809, when fused to a human IgG1 Fc domain, showed transmission reduction of up to 47% at a concentration of 100 µg/mL. While nanobody W2809 has a larger epitope footprint compared to nanobodies W2810 and W2812, which bind to Pfs230 D13, nanobody W2809 has the lowest affinity to Pfs230 D13D14, with a *K_D_* of 46.6 nM. Future nanobody engineering approaches using information from our high-resolution crystal structures may yield a higher affinity nanobody to Pfs230 D13D14, which could improve its transmission-blocking potency. Nanobody W2811 showed up to 44% reduction in transmission when assessed by SMFA. Nanobody W2811 and W2812 bind in the same epitope bin and have similar affinities for Pfs230 D13 but W2812 has negligible transmission-blocking capabilities. To improve the potency of nanobody W2811, understanding its precise epitope using X-ray crystallography will be required. Our successful expression of well-folded recombinant Pfs230 D13D14, as evidenced by our crystal structures of these domains, will facilitate the isolation of a larger diversity of antibodies using single B cell approaches for the isolation of human monoclonal antibodies from naturally infected *P. falciparum* individuals.

We show that TB31F can disrupt complex formation between Pfs230 and Pfs48/45. TB31F is one of the most potent anti-Pfs48/45 mAbs, showing 80% reduction in transmission at a concentration of 0.5-1.0 µg/mL^53^, and is the humanized version of mAb 85RF45.1, which reduces transmission by 80% at a concentration of 1-2 μg/mL^53,65,72,73^. A recent Phase 1 clinical trial in malaria naïve individuals showed TB31F to be well-tolerated and capable of blocking *P. falciparum* transmission from humans to mosquitoes^74^. Overlaying the epitope footprint of TB31F with our cryo-EM of Pfs230-Pfs48/45, we show that this mAb directly interferes with Pfs230 binding to Pfs48/45. The crystal structures of TB31F in complex with Pfs48/45 D3 suggest that its epitope is relatively conserved^53,65^ and reported field polymorphisms do not substantially affect antibody binding or transmission-reducing activity^31,53,65,73,75^. Based on our cryo-EM structure, we predict that anti-Pfs48/45 D3 mAbs 32F3, RUPA-29 and RUPA-47^28,31^ could also directly interfere with Pfs230 binding, but this would require future validation.

We show that a *P. falciparum* transgenic line that does not express D13D14 has a strong defect in surface localization of Pfs230 in activated male and female gametes compared to the parental line. Using independent clones of Δ230D1314, we show that over 90% of activated female gametes have no Pfs230 localization on their surfaces compared to 14% in NF54/iGP2. This phenotype is similar to the absence of Pfs230 on activated gametes observed for a Pfs48/45 knockout line^20^. Previous studies have also shown that 11.9% of the activated macrogametes showed only Pfs230 staining on the membrane surface without Pfs48/45 staining, which is similar to our results^43^. At present we do not know what is responsible for the 12 to 14% of residual membrane localization of Pfs230 in the absence of Pfs48/45 and in the Δ230D1314 line. One possibility is that in the absence of Pfs48/45, Pfs230 may remain associated with the gamete surface through its interactions with the PfCCp multimeric protein complex which is anchored by Pfs25^76^, but without the main anchoring mechanism through Pfs48/45, Pfs230 is unstable and dissociates from membrane. Pfs230 expression is not affected by Pfs48/45 disruption as determined by western blotting^26^ and interestingly, in both the 230Δ1314 line generated in this study and the previously described Pfs48/45 knockout line^20^, Pfs230 is still expressed in stage V gametocytes. We propose that this localization represents secreted Pfs230 and its accumulation below the parasitophorous vacuolar membrane. However, when gametocytes are activated, the macrogametes and microgametes egress from the red blood cell and the anchoring of Pfs230 to the gamete membrane via its interaction with Pfs48/45^20–22^ becomes essential for successful surface localization. In addition, 230Δ1314 parasites are defective in parasite fertilization as observed by the significant decrease in oocyst development in the mosquito midgut. These results show that D13 and D14 of Pfs230 are critical for its function and provides a novel immunogen for transmission blocking vaccines.

Our Pfs230 D13D14 mRNA-LNP immunogens at 10 μg per immunization dose were able to elicit antibodies specific to these Pfs230 domains. However, only immunization with 230D13D14S mRNA-LNP generated antibodies that potently reduce transmission up to 98%. Currently, we do not fully understand why 230D13D14L mRNA-LNP immunization does not generate transmission blocking antibodies. We hypothesize that the expressed protein may be unstable and degrading but this will require future work to determine the exact reason. At present, our mRNA designs only included a signal peptide and not any membrane anchor motifs. A recent study on Pfs25 and Pfs230D1 showed mRNA constructs with a GPI anchor or transmembrane domain generated sera with higher transmission-blocking activity^77^. Next generation designs for Pfs230 D13D14 should include a signal peptide and either a GPI anchor or transmembrane domain to test whether membrane anchorage also elevates antibody titers using our vaccines. In addition, it will be important to compare which mRNA-LNP vaccines of Pfs230D1, Pfs230 D13D14, Pfs230D13 and Pfs230D14 can elicit the highest potency in transmission-blocking. Furthermore, there has not been any studies using mRNA-LNP delivery of Pfs48/45 as a transmission-blocking vaccine. Future studies should examine if a combination mRNA-LNP vaccines with Pfs48/45 and the most potent fragment of Pfs230 will provide a higher transmission-reducing functional activity compared to individual subunit vaccines.

## Methods

### Generation of transgenic Pfs230FL and 230Δ1314 3xFLAG-TwinStrepII-tagged parasites

To introduce 3xFLAG-TwinStrepII tag to the C-terminal end of Pfs230, we used the NABS4_3xFLAG-TwinStrepII_5’-3’_Pfs230 construct. This construct was generated as follows: 3xFLAG-TwinStrepII was synthesized as a gBlock (Integrated DNA Technologies) and cloned into the NABS4 vector using XhoI/XmaI. A 5’ homology region (HR1) of Pfs230 followed by a recodonized coding sequence was cloned into the NABS4_3xFLAG-TwinStrepII plasmid using NotI and XhoI restriction sites. Subsequently the 3’ homology region (HR2) for targeting Pfs230 was cloned in using KpnI and PstI sites.

A Cas9/guide RNA expressing plasmid^78^ with guide sequence targeting Pfs230 (pUF1-Cas9_Pfs230_g8947) was transfected with NotI and PstI linearized NABS4_3xFLAG-TwinStrepII_5’-3’_Pfs230 into NF54/iGP2 parasites^54^, using the Lonza Nucleofector Transfection 2b device and Basic Parasite Nucleofector Kit 2 as previously described^79^. In brief, synchronized schizonts were purified using a Percoll (Cytiva) density gradient. 100 µg each of the linearized repair template and Cas9/guide expressing plasmid pUF1-Cas9_Pfs230_g8947 was resuspended in 100 µL of supplemented Basic Parasite Nucleofector Solution 2. 50 μL of packed schizonts were resuspended in the DNA/Nucleofector solution and transferred to the cuvette for electroporation (program U-33). Electroporated parasites were transferred to a prewarmed 1.5 mL Eppendorf tube using 500 μL of culture medium and 50 μL of fresh red blood cells and incubated at 37 °C at 650 RPM for 30 mins. Transfected parasites were returned to regular culture conditions. The following day, 2.5 nM of the selection drug WR99210 (Jacobus Pharmaceutical Co) was added and maintained until >2% parasitemia was observed.

To generate the 230Δ1314 line, NABS4_3xFLAG-TwinStrepII_5’-3’_Pfs230 was modified in the following way. The nucleotide sequence 5’ of D13 of Pfs230 was PCR amplified to use as the 5’ homology region (HR3) and cloned into the vector using NotI and XhoI sites, and this plasmid is called NABS4-3xFLAG TwinStrepII_Pfs230_3’_D13D14_5’. Parasite transfection with this construct was performed as described above with the same Cas9/guide RNA expressing plasmid pUF1-Cas9_Pfs230_g8947. Clonal parasite populations were obtained using limited dilution cloning. Parasites were assessed for integration of the 3xFLAG-TwinStrepII tag at the 3’ of the gene by extracting gDNA from 0.15% saponin lysed parasite material (DNeasyBlood and Tissue Kit, Qiagen) and using genotyping PCR (CloneAmp HiFi PCR premix, Takara). In addition, western blotting was used to confirm the expression of the 3xFLAG-TwinStrepII tag on Pfs230 (Figure S1C). All primer and synthetic DNA sequences are provided in Table S4.

### Asexual blood stage growth and gametocyte development

Asexual *P. falciparum* parasite cultures were maintained with O^+^ red blood cells at 4% haematocrit in RPMI 1640, GlutaMAX, HEPES (Thermo Fisher) supplemented with: hypoxanthine (12.5 mg/L) (Sigma), D-glucose (2 g/L) (Thermo Fisher), gentamycin (20 mg/L) (Thermo Fisher), 0.25% (w/v) AlbuMAX II (Thermo Fisher), and 5% (v/v) O^+^ human serum (Australian Red Cross, Lifeblood). Cultures were maintained by routine sub-culturing and media replacement as necessary, and NF54/iGP2 maintained as asexual forms through the addition of 2.5 mM D-(+)-glucosamine hydrochloride (Sigma). Gametocytes were induced as previously described for the NF54/iGP2 line^54^ and when the culture was majority stage V gametocytes, they were maintained at 37°C until red blood cells were lysed using 0.03% saponin. Parasites were washed in ice cold phosphate buffered saline (PBS) supplemented with cOmplete protease inhibitor (Roche) and stored at −80°C until protein purification.

### Purification of endogenous Pfs230-Pfs48/45 complex from stage V gametocytes

Parasite pellets were resuspended in lysis buffer, comprising 50 mM HEPES pH 8.0, 150 mM NaCl, 10 mM EDTA, 10% (v/v) glycerol, and either 1% N-dodecyl β-D-maltoside (DDM) or glyco-diosgenin (GDN), and incubated at room temperature (RT) for 45 min on rollers. The soluble lysate was separated from the insoluble fraction by centrifuging at 16,000 *x* g for 1 h at 4°C and incubated with StrepTactin XT resin (IBA Lifesciences, equilibrated in lysis buffer) for 2 h on rollers at 4°C. The resin was transferred to a gravity chromatography column and washed with wash buffer containing reduced detergent concentrations. Elution was performed with 1x Buffer BXT (IBA Lifesciences) containing 5% (v/v) glycerol and 0.02% DDM or 0.0075% GDN in addition to 100 mM Tris pH 8.0, 1 mM EDTA and 50 mM biotin. Fractions containing the proteins of interest were pooled and concentrated using a 30 kDa molecular weight cutoff (MWCO) Millipore concentrator. Size exclusion chromatography (SEC) was used to assess sample quality and as a last purification step. The sample was applied to a Superose 6 3.2/300 column (Cytiva) equilibrated in 50 mM HEPES pH 8.0, 150 mM NaCl, 3% (v/v) glycerol, 0.02% DDM or 0.0075% GDN. Samples from fractions of interest, alongside 1 µg SARS-CoV-2 Spike^59^, were loaded onto a 3-8% NuPAGE Tris-Acetate gel (Invitrogen) and run in 1X Tris-Acetate running buffer for 1 h at 150 V. Gels were stained with InstantBlue Coomassie protein stain to identify fractions containing the complex. Protein in DDM-containing buffer was used for cryo-EM and the protein in GDN for crosslinking-mass spectrometry approaches.

### Purification of recombinant Pfs230 domains

Codon optimized (*Spodoptera frugiperda, Sf*) DNA corresponding to amino acids K2828-E3111 of Pfs230 (D13D14), D2446-T2818 of Pfs230 (D11D12), T1281-G1549 of Pfs230 (D5D6) and N2450-S2731 of Pvs230 (PvD13D14) were cloned into a modified form of baculovirus transfer vector pAcGP67-A. The Pfs230 sequence is in frame with the GP67-signal sequence and an N-terminal octa-histidine tag followed by a Tobacco Etch Virus (TEV) protease cleavage site. The recombinant protein was produced using *Sf*21 cells (Life Technologies) cultured at 28°C in Insect-XPRESS Protein-free Insect Cell Medium supplemented with L-glutamine (Lonza). A cell culture of ∼1.5 - 2.0 x 10^6^ cells/mL was inoculated with the third passage stock of virus and incubated for three days at 28°C. Cells were separated from the supernatant by centrifugation at 17,000 *x* g for 30 min. The supernatant was sterile filtered with 0.45 μm filters and concentrated via tangential flow filtration using a 10 kDa MWCO cassette (Millipore). The concentrated supernatant was sterile filtered with a 0.45 μm filter and dialyzed into 30 mM Tris-HCl pH 7.5, 300 mM NaCl. The dialyzed sample was incubated with Ni-NTA resin (Qiagen) for 1h at 4 °C on a roller shaker. Ni-IMAC using a gravity flow chromatography column was performed. The resin was washed with 10-20 column volumes of 30 mM Tris-HCl pH 7.5, 300 mM NaCl, followed by further washes with stepwise increases in imidazole concentration. TEV protease was added to the pooled fractions containing Pfs230 D13D14 while dialyzing into 30 mM Tris-HCl pH 7.5, 300 mM NaCl to remove the N-terminal tag. The solution was incubated with Ni-NTA resin (Qiagen) for 1h at 4°C on a roller shaker before untagged Pfs230 D13D14 was separated from His-tagged TEV protease and un-cleaved protein via Ni-IMAC purification. The flow-through was concentrated and applied to a Superdex 200 Increase 10/300 SEC column (Cytiva) pre-equilibrated with 20 mM HEPES pH 7.5, 150 mM NaCl.

### Cryo-EM sample preparation and data acquisition

Cryo-EM grids were prepared using a multiple blotting approach to concentrate the particles in the holes. The main SEC peak fraction was used and 3.5 μL of sample was applied to a plasma-cleaned R1.2/1.3 UltrAuFoil grid, followed by blotting of the grid at a force of 3 N for 4 s. Four to six applications were performed, with the last blotting performed for 7 s before plunge freezing the grid in liquid ethane using a Vitrobot Mark IV (ThermoFisher) at 20 °C and 100 % humidity. Cryo-EM grids were screened on the Arctica and subsequently imaged on a Titan Krios microscope at the Ian Holmes Imaging Centre, Bio21 (IHIC, Melbourne) operated at 300 kV using a Falcon 4 detector in electron counting mode. A total of 35,375 movies were collected from four grids at a pixel size of 0.808 Å/pixel, with a total dose of 50 e /Å^2^. Images were acquired using a 70 μm objective aperture and a defocus range of −0.4 to −1.6 μm.

### Data processing

Movies were motion corrected and contrast transfer function parameters estimated in cryoSPARC v4.6.0^80^ (Figure S3 and Figure S4). Datasets from each grid were pre-processed individually. Approximately 100 particles were manually picked from each set of micrographs before blob tuner picking in cryoSPARC. The picked particles were extracted (360 pixels, 2x binned to 180 pixels) and subjected to two-dimensional (2D) classification separately. Several rounds of 2D-classification were performed and particles from poorly defined 2D-classes iteratively excluded. About 20,563 particles of grid 1 were used for *ab initio* reconstruction into four classes. Particles of grids 1 and 2 were combined and subjected to homogenous refinement, further rounds of 2D-classification and heterogeneous refinement, which separated the particles into intact Pfs230-Pfs48/45 complex (53,851 particles) and junk (19,570 particles). Particles of grid 3 and 4 (97,225 particles) were added and further rounds of 2D-classification and heterogenous refinement used to clean up the particle stack. Particles were re-extracted (360 pixels) and subjected to local per-particle contrast transfer function (CTF) refinement. Following heterogenous refinement, a final non-uniform refinement with the final dataset of 87,061 particles yielded a consensus map of 3.36 Å based on the gold standard Fourier shell correlation (FSC) cutoff of 0.143.

To improve the density of the Pfs230 domains D1-D8, the map was subjected to a focused refinement, resulting in an overall resolution of 4.86 Å. Our consensus and local refinement cryo-EM maps of the Pfs230-Pfs48/45 complex showed preferred orientation bias, therefore, to aid in model building, we used EMReady (v2.0)^81^ on unsharpened maps for better interpretation of the maps. We tried different values for the flag “Stride”, which controls the step of the sliding window for cutting the input map into overlapping boxes that range from 12 to 48 but found the default setting of 12 worked the best. The interpretability of the consensus cryo-EM map improved for the region spanning Pfs230 D9-14 and Pfs48/45, allowing confident tracing of the backbone in most regions.

A composite map was generated from the EMReady processed consensus map and the local refinement map comprising Pfs230 D1-D8. Local resolution of the maps was estimated in cryoSPARC (v4.6.0)^80^. Due to the observed variations in local resolution and preferred orientation, we report only the global positions of individual domains that could be modelled with high confidence.

### Model building and refinement

AlphaFold2^82^ and AlphaFold2 Multimer (v2.3.2)^83^ predicted models were calculated using the WEHI high-performance computing system. The highest scoring model of the Pfs230-Pfs48/45 complex was used for model building into the composite map. The model was divided into three regions. Regions spanning Pfs230 D1-D6 (residues 527-1683), Pfs230 D7D8 (residues 1684-2037) and Pfs230 D9-D14 (residues 2038-3133)-Pfs48/45 (residues 28-428) were rigidly fitted independently into the map using ChimeraX (v1.8)^84^. Namdinator^85^, a web-based automated molecular dynamics based flexible fitting program, was used to improve initial fitting of the models. The Pfs230 D1-D8 model was in the lower resolution part of the map; therefore, all side chains were truncated and no further fitting was performed, except rigid body fitting in Phenix (v1.21.1)^86^ to generate the model-map statistics (Table S1). The Pfs230 D9-D14-Pfs48/45 model was in the higher resolution part of the map. To further improve the fitting, an iterative refinement in Coot^87^ and Phenix (with global minimization and secondary structure restraints) was performed. Since our cryoEM maps have regions of preferred orientation, we only fitted the global orientation of the domains that we can model confidently. Crystal structures of the Pfs48/45 ectodomain (PDB IDs 7ZXG and 7ZXF) were used to aid model building of its domains D2 and D3 in the map.

### Crosslinking mass spectrometry sample preparation and analysis

Sulfosuccinimidyl 4,4’-azipentanoate (Sulfo-SDA) (Thermo Scientific Pierce) was dissolved freshly in 50 mM HEPES pH 8.0, 150 mM NaCl, 5% (v/v) glycerol, 0.0075% GDN to a concentration of 1 mg/mL. Crosslinking reactions were performed with a two-fold dilution series of Sulfo-SDA ranging from 1.0 mg/mL to 0.06 mg/mL. Purified Pfs230-Pfs48/45 complex (5 μL, ∼0.1 mg/mL, ∼1.25 μM) was mixed with 5 μL Sulfo-SDA and incubated in the dark at RT for 30 min. To activate Sulfo-SDA, the samples were irradiated with an UV mercury-xenon arc lamp at 1000 watts on ice for 1 min. Crosslinked samples were analyzed on a 3-8% NuPAGE Tris-Acetate gel (Invitrogen). Protein without crosslinker and protein mixed with the highest crosslinker concentration, which lacked UV exposure, were used as controls. Gel bands corresponding to the crosslinked Pfs230-Pfs48/45 heterodimer were excised and enzymatically digested with trypsin as previously described^88^.

Reconstituted peptides were analyzed on an Orbitrap Eclipse™ Tribrid mass spectrometer that is interfaced with a Vanquish Neo liquid chromatography system. Samples were loaded onto a C18 fused silica column (inner diameter 75 µm, OD 360 µm × 15 cm length, 1.6 µm C18 beads) packed into an emitter tip (IonOpticks) using pressure-controlled loading with a maximum pressure of 1,500 bar, which is interfaced to an Orbitrap Eclipse™ Tribrid (Thermo Scientific) mass spectrometer using EASY-nLC source and electrosprayed directly into the mass spectrometer. Peptides were eluted using a 120-minute linear gradient of 3% to 30% of acetonitrile into the mass spectrometer operating in data-dependent mode (scan parameters detailed in Data S1).

Raw data files were converted to MGF files using msConvert^89^. MGF files were searched against a fasta file containing the Pfs230 and Pfs48/45 sequences using xiSEARCH software (version 1.7.6.7)^90^ with the following settings: crosslinker = multiple, SDA and noncovalent; fixed modifications = Carbamidomethylation (C); variable modifications = oxidation (M), sda-loop (KSTY) DELTAMASS:82.04186484, sda-hydro (KSTY) DELTAMASS:100.052430; MS1 tolerance = 6.0ppm, MS2 tolerance = 20.0ppm; losses = H_2_O,NH_3_, CH_3_SOH, CleavableCrossLinkerPeptide:MASS:82.04186484). False discovery rate estimation was performed with the in-built xiFDR set to 5%. Data were visualized using the xiVIEW software^91^.

### Immunofluorescence assay on *Plasmodium falciparum* parasites

NF54/iGP2 stage V gametocytes were cultured as described above. Thin blood smears were made on glass slides, fixed in equal parts of ice-cold acetone and methanol for 10 min and allowed to air dry. The slides were incubated with 3% (w/v) bovine serum albumin (BSA)/PBS for 1 h then washed three times in 1x PBS prior to addition of the primary antibodies. The primary antibodies or nanobodies were diluted in 3% BSA/PBS. Anti-Pfs230 LMIV230-01 directly conjugated to Alexa Fluor 555 was used at a final concentration of 7.5 to 10 μg/ml, anti-Pfs48/45 TB31F directly conjugated to Alexa Fluor 647 at final concentration 7.5 to 10 μg/mL, the Pfs230 D13D14 nanobodies W2809, W2810, W2811 and W2812 and the negative control SARS-CoV-2 nanobody WNb7 were used at 4 μg/mL^59^. Primary antibodies were incubated on the slide for 2 h within a humidified chamber. After washing as described, goat anti-alpaca IgG VHH domain conjugated to Alexa Fluor 488 (Jackson ImmunoResearch) (diluted 1:1000 in 3% BSA/PBS) was used as a secondary antibody and incubated for 2 h within a humidified chamber. For the mouse serum IFA, pooled serum was diluted to 1:500 and donkey anti-mouse conjugated to Alexa Fluor 647 (Thermo Scientific) diluted 1:1000 in 3% BSA/PBS was used as a secondary antibody. The slides were mounted with a coverslip using VECTASHIELD PLUS antifade mounting medium with DAPI (Vector Laboratories) and sealed with nail polish. Images were acquired on a Zeiss LSM 980 using confocal mode with a 63x (1.4NA) objective lens with oil immersion using the 405, 488 and 561 lasers. Images were processed using FIJI ImageJ software (v2.14.0/1.54)^92^ and the Z-projection max intensity images were used in the figures.

### Surface immunofluorescence assay of *P. falciparum* activated gametes

Stage V gametocytes were purified from uninfected red blood cells using a Percoll (Cytiva) centrifugation density gradient. Gamete activation was induced by incubation in ookinete medium (RPMI with 100 μM xanthurenic acid)^93^ at RT for 30 min. To obtain macrogamete images, gametes were washed 3X with 0.5% fetal bovine serum (FBS), 0.05% sodium azide in PBS prior to incubation with anti-Pfs48/45 TB31F directly conjugated to Alexa Fluor 647 and anti-Pfs230 LMIV230-01 directly conjugated to Alexa Fluor 555. Antibody incubation was performed at 4°C in the dark with gentle shaking for 1 h. Activated gametes were washed as described above, then fixed by resuspension in 2% formaldehyde in PBS for 20 min. A final series of washes was performed prior to adherence of gametes to a 0.1 mg/mL erythroagglutinin PHA-E (Sigma-Aldrich) coated coverslip. Slides were then mounted and imaged as described above using the 405, 561 and 639 lasers. Microgamete SIFA was performed by resuspending stage V gametocytes in ookinete medium supplemented with the Alexa Fluor conjugated antibodies. The activating gametes were then allowed to settle onto coverslip, incubated in a humidified chamber in the dark for 30 minutes, then fixed, mounted, and imaged as described above. Z-projection images were generated in FIJI ImageJ and brightness and contrast were adjusted for visualization. Automated quantitation of SIFA of macrogametes were performed as follows. Cell segmentation was performed via a custom trained cellpose^94,95^ network on maximum intensity projected brightfield images. Cell scoring was performed via a custom FIJI^92^ macro with manual thresholds set for mean intensity of maximum intensity projected images of 647 and 555 channels. Mask images showing the segmentation and scoring of each cell were produced for manual validation.

### Western blotting

Parasite lysate was resuspended in either reducing or non-reducing sample buffer and separated on a 3 – 8% NuPAGE Tris-Acetate SDS-PAGE (ThermoFisher Scientific). Samples were transferred onto PVDF membranes (iBlot™ 2 Transfer Stacks, ThermoFisher Scientific) and blocked overnight in 10% skim milk in PBST. Blots were probed with anti-Pfs230 LMIV-230-01 and anti-FLAG followed by the appropriate secondary antibodies conjugated to HRP, and anti-Strep-HRP. Chemiluminescence was detected using SuperSignal West Pico PLUS Chemiluminescent Substrate (ThermoFisher Scientific). The HRP signal was inactivated using 0.025% sodium azide in PBST and re-probed with anti-HSP-70 followed by appropriate secondary and imaged as above.

### Alpaca immunization and isolation of Pfs230 D13D14 nanobodies

One alpaca was immunized six times with approximately 200 μg of recombinant Pfs230 D13D14 protein on days 0, 14, 21, 28, 35 and 42. The adjuvant used was GERBU FAMA. Immunization and handling of the alpacas for scientific purposes was approved by Agriculture Victoria, Wildlife & Small Institutions Animal Ethics Committee, project approval No. 26-17. Blood was collected three days after the last immunization for the preparation of lymphocytes. Nanobody library construction was carried out according to established methods as described^96^. Briefly, alpaca lymphocyte mRNA was extracted and amplified by RT-PCR with specific primers to generate a cDNA nanobody library. The library was cloned into a pMES4 phagemid vector containing 10^4^ unique nanobodies, amplified in *E. coli* TG1 strain and subsequently infected with M13K07 helper phage for recombinant phage expression. Biopanning for recombinant Pfs230 D13D14 nanobodies using phage display was performed as previously described with the following modifications^59^. Phages displaying Pfs230 D13D14-specific nanobodies were enriched after two rounds of biopanning with 1 μg of immobilized Pfs230 D13D14 complex. After the second round of panning, we screened 94 clones from round two by ELISA and selected positive hits for further analysis. Positive clones were sequenced and annotated using PipeBio. Positive clones were sequenced and annotated using PipeBio. Distinct nanobody clonal groups were identified based on differences in the amino acid sequence of the complementary determining region 3 (CDR3) that vary by at least one amino acid.

### Purification of nanobodies and Fc-tagged nanobodies

Nanobodies were expressed in *Escherichia coli* WK6 cells and purified as previously described^46^. Briefly, bacteria were induced at an OD_600_ of 0.7 with 1 mM IPTG and grown overnight at 28°C. Cell pellets were harvested, resuspended in 20% (w/v) sucrose, 20 mM imidazole pH 7.5, 150 mM NaCl PBS and incubated for 15 min on ice. 5 mM EDTA pH 8.0 was added followed by an incubation on ice for 20 min. After this incubation, 10 mM MgCl_2_ was added, and periplasmic extracts were harvested by centrifugation. The supernatant was loaded onto a 1 mL HisTrap Excel column (GE Healthcare), washed and the nanobodies eluted with 400 mM imidazole pH 7.5, PBS, before being concentrated and buffer exchanged into PBS.

Nanobody sequences of W2809, W2810, W2811 and W2812 were subcloned into a derivative of pHLSec containing the hinge and Fc region of human IgG1 to produce nanobody-Fc tagged fusion proteins in Expi293 cells via transient transfection. Restriction-ligation cloning with PstI and BstEII restriction enzymes was used to introduce the respective sequences. The supernatant was harvested six days after transfection and applied to 1 mL HiTrap Mab Select PrismA affinity columns (Cytiva). Nanobody-Fcs were eluted in 100 mM citric acid pH 3.0, neutralized by the addition of 1 M Tris-HCl pH 9.0 and subsequently buffer exchanged into PBS.

### Next-generation sequencing and analyses of nanobody phage libraries

Plasmids from phage libraries were digested and gel purified to isolate the nanobody domain sequences. Paired-end 2×300 bp sequencing libraries were prepared using the NEBNext Multiplex Oligos for Illumina (Cat # E7395) in a PCR-free manner according to the manufacturer’s instructions. The samples were size-selected using AMPure XP beads, quantified by qPCR using the KAPA Library Quantification Kit and sequenced on an Illumina NextSeq 2000 instrument. Raw sequencing reads were trimmed to remove sequencing adapters with TrimGalore v0.6.7 (https://github.com/FelixKrueger/TrimGalore) and then merged using FLASH v1.2.11^97^. Annotation of nanobody sequences was performed using IgBLAST v1.19.0^98^ with an alpaca reference database built from IMGT^99^. Nanobody sequences were collapsed into clones at the CDR3 level, and the counts of clones were normalized through conversion to counts per million (CPM). Cladograms were generated from generalized Levenshtein distances between CDR3s with the Neighbor-Joining method^100^ from the phangorn (v2.12.1)^101^ package and plotted using the ggtree (v3.14.0)^102^ package in R (v4.4.1)^103^. Amino acid logos were made using the ggseqlogo R package (v0.2.0)^104^.

### ELISA for nanobody binding

The 96-well flat-bottomed MaxiSorp plates were coated with 125 nM of recombinant protein, as indicated, in 50 μL PBS at RT for 1 h. All washes were done three times using PBS and 0.05% (v/v) Tween-20 (PBST), and all incubations were performed for 1 h at RT. Coated plates were washed and blocked by incubation with 10% (w/v) skim milk solution. Plates were washed and then incubated with nanobody at a concentration of 125 nM. The plates were washed and incubated with mouse anti-His (Bio-Rad MCA-1396; 1:1000) followed by horseradish peroxidase–conjugated goat anti-mouse secondary antibody (MerckMillipore AP124P, 1:1000). After a final wash, 50 μL azino-bis-3-ethylbenthiazoline-6-sulfonic acid (ABTS liquid substrate; Sigma 11684302001) was added and incubated in the dark at RT, and 50 μL 1% (w/v) sodium dodecyl sulfate (SDS) was used to stop the reaction. Absorbance was read at 405 nm, and all samples were done in duplicate.

### Affinity determination, epitope binning and domain specificity using biolayer interferometry (BLI)

Affinity determination measurements were performed on the Octet RED96e (FortéBio). Assays using monomeric nanobodies were performed using Penta-His antibody capture sensor tips (Octet®HIS1K). For assays using nanobody-Fcs, anti-hIgG Fc capture sensor tips (Octet^®^ AHC) were used. All measurements were performed in kinetics buffer (DPBS pH 7.4 supplemented with 0.1% (w/v) BSA and 0.05% (v/v) Tween-20) at 25°C. After a 60 s baseline step, nanobodies or nanobody-Fcs (5 μg/mL) were loaded onto sensors until a response of 0.5 nm, followed by a 60 s baseline step. Association measurements using nanobodies were performed using a two-fold dilution series of untagged Pfs230 D13D14 from 6 to 200 nM for 180 s for determination of affinity to D13D14 and a single 100 nM concentration of Pfs230 domains to determine domain specificity of nanobodies. Dissociation was measured in kinetics buffer for 180 s. Sensor tips were regenerated using five cycles of 5 s in 100 mM glycine pH 1.5 and 5 s in kinetics buffer for both HIS1K-sensors and AHC-sensors. Baseline drift was corrected by subtracting the response of a nanobody-loaded sensor incubated in kinetics buffer only. Curve fitting analysis was performed with Octet Data Analysis 10.0 software using a global fit 1:1 model to determine *K*_D_ values and kinetic parameters. The mean kinetic constants reported are the result of two to three independent experiments. Our data shows R^2^ values of > 0.998 and Ξ^2^ < 0.03 representing a statistically reliable goodness-of-fit of experimental data against the expected binding model (Dataset S2).

For epitope binning experiments, 50 nM untagged Pfs230 D13D14 was pre-incubated with each nanobody at a 10-fold molar excess for 1 h at RT. A 30 s baseline step was established between each step of the assay. The NTA sensors were first loaded with 10 μg/mL of nanobody for 5 min. The sensor surface was then quenched by dipping into 10 μg/mL of an irrelevant nanobody for 5 min. Nanobody-loaded sensors were then dipped into premixed solutions of Pfs230 D13D14 and nanobody for 5 min. Nanobody-loaded sensors were also dipped into Pfs230 D13D14 alone to determine the level of Pfs230 D13D14 binding to immobilized nanobody in the absence of other nanobodies. Percentage competition was calculated by dividing the max response of the premixed Pfs230 D13D14 and nanobody solution binding by the max response of Pfs230 binding alone, multiplied by 100.

### Pull down assay of endogenous Pfs230-Pfs48/45 complex for blocking nanobodies

Parasite lysates for the pull down assay was prepared by lysing frozen saponin-treated parasite pellets in 50 mM HEPES pH 8.0, 150 mM NaCl, 10 mM EDTA, 10% (v/v) glycerol, 1% DDM, followed by centrifugation to obtain soluble lysate. Antibody or nanobody-Fc was added to a final concentration of 1.3 μM to 15 μL of parasite lysate in a 50 μL reaction volume with assay buffer (50 mM HEPES pH 8.0, 150 mM NaCl, 3% (v/v) glycerol, 0.02% DDM), and Input samples were taken. Input and Unbound samples were comprised of 5 μL sample and 5 μL of 2x SDS-PAGE non-reducing sample buffer. 10 μL of a 50% slurry of StrepTactin XT resin (IBA Lifesciences) in assay buffer was added and incubated for 2 h on rollers at 4°C. Samples were centrifuged at 1500 *x* g for 2 min to pellet the resin, and Unbound samples were taken. The resin was washed three times in assay buffer by centrifugation at 1500 *x* g for 2 min. Proteins were eluted from the resin by boiling for 5 min in 15 μL of 2x SDS-PAGE non-reducing sample buffer, followed by centrifugation at 1500 *x* g for 2 min, and 10 μL was taken for the Eluate sample. 10 μL each of Input, Unbound and Eluate samples were run on 3-8% NuPAGE Tris-Acetate gels (ThermoFisher Scientific) in 1X NuPAGE Tris-Acetate running buffer at 150 V for 60 min, then transferred onto a PVDF membrane. All antibody incubations were performed for 1 h at RT, followed by three washes with PBS 0.1% (v/v) Tween-20 at RT for 5 min. The blot was blocked with 10% (v/v) milk in PBS 0.1% Tween-20 overnight at 4°C, then washed and incubated with 2 μg/mL of anti-Pfs48/45 mouse monoclonal antibody 3E12^29,43^. After washing, the blot was incubated with HRP-conjugated goat anti-mouse antibody (MerckMillipore AP124P, 1:1000). Finally, the blot was incubated with HRP-conjugated NWSHPQFEK rabbit polyclonal antibody (ThermoFisher Scientific A00875-40, 1:1000). After a final wash with PBS for 10 min, bands were detected using SuperSignal West Pico PLUS Chemiluminescent Substrate (ThermoFisher Scientific) and imaged using a ChemiDoc Imaging System (Bio-Rad).

### Crystallization and structure determination of Pfs230 D13D14-nanobody complexes

For crystallization, stable complexes were formed between Pfs230 D13D14 and monomeric nanobodies by mixing antigen and nanobody at a molar ratio of 1:1.5 and incubating for 1 h on ice. Pfs230-nanobody complexes were purified from excess nanobody using SEC in 20 mM HEPES pH 7.5, 150 mM NaCl. Sitting drop vapor diffusion crystallization trials were performed with 5 and 12 mg/mL of complex. Initial crystals of Pfs230 D13D14-W2809 grew in 10% polyethylene glycol (PEG) 8000, 0.2 M sodium acetate, 0.1 M imidazole pH 8.0 at 20°C and were optimized using hanging drop vapor diffusion by increasing the PEG concentration to 12% as well as incorporating low percentages of cryoprotectant (4% glycerol) within the mother liquor. Crystals of Pfs230 D13D14-W2812 grew in 5% PEG 8000, 5% PEG 10000, 5% PEG 6000, 0.15M ammonium acetate, 0.1 M sodium citrate pH 5.0 at 4 °C and Pfs230 D13D14-W2810 crystals in 1.4 M sodium malonate dibasic monohydrate pH 7.0 at 20 °C. For cryo-protection, a premix of 30% PEG 8000 and 10% glycerol within the mother liquor was used for Pfs230 D13D14-W2809 crystals, 30% glycerol in mother liquor was used for Pfs230 D13D14-W2812 and 30% ethylene glycol in mother liquor for Pfs230 D13D14-W2810 crystals, respectively.

Data sets were collected from single crystals on the MX2 beamline at the Australian Synchrotron and processed using the XDS package^105^. Phaser^106^ in the CCP4 suite^107^ was used for molecular replacement using coordinates of a known nanobody structure (PDB ID 8AOK) and an AlphaFold2 (v.2.3.2) predicted model for Pfs230 D13D14 as search models. Structure refinement was undertaken through iterative rounds of model building in Coot and refinement in Phenix. Figures were prepared in ChimeraX^84^. Interfaces and interactions were analyzed and buried surface areas calculated using PISA^108^. The three structures have been deposited in the PDB under accession codes 9E7N (Pfs230 D13D14-W2809), 9E7O (Pfs230 D13D14-W2810) and 9E7P (Pfs230 D13D14-W2812).

### Standard membrane feeding assays

Standard membrane feeding assays were conducted at Seattle Children’s Research Institute using the *P. falciparum* NF54 line^109^. To initiate gametocyte development, healthy asexual cultures, primarily containing ring-stage parasites, were set up at 1% parasitemia and 5% hematocrit in a 6-well plate. Gametocyte cultures were maintained for 14 days, with daily media changes. On day 15, stage V gametocytes were evaluated, and female *Anopheles stephensi* mosquitoes (4-7 days old) were fed on mature gametocyte cultures at 0.3% gametocytemia and 50% hematocrit. Nanobody-Fcs were added to the blood meal at a final concentration of 100–200 µg/mL. LMIV230-01, used as a positive control, was included at a concentration of 200 µg/mL. After one week, midguts were dissected, and oocyst numbers were enumerated. Standard membrane feeding assays for the Δ230D1314 lines were conducted at WEHI using a similar approach as above without the addition of nanobodies. Assays performed using pooled mouse pre-immune and 230D13D14S and 230D13D14L mRNA-LNP sera were conducted at WEHI using a similar approach as above with pooled mouse serum added at a final concentration of 1:5.

### Papua New Guinea (PNG) study populations

Plasma samples were obtained from children (aged 5-10 years) in PNG that participated in a randomized double-blind drug trial^110^. For this experiment, the final time point of the sample collection at 36 weeks post-enrolment was used. Thirty-two samples were selected from children who were currently infected with *P. falciparum* at this timepoint; of these, 13 were co-infected with *P. vivax*. Forty samples were included from PNG children who did not have a current *Plasmodium* infection at the time of sample collection, but live in the same endemic area, with 27.5% having had PCR-detectable *P. falciparum* infections during the prior 36-week period (47.5% had PCR-detectable *P. vivax* infections). These children were randomly selected and had equivalent demographics to the 32 *P. falciparum* positive children. Forty negative control samples were selected from the Volunteer Biospecimen Donor Registry (VBDR) in Melbourne. The studies involving human participants were reviewed and approved by the Walter and Eliza Hall Institute Human Research Ethics Committee (13/07), the PNG Institute of Medical Research Institutional Review Board (0908) and the PNG Medical Advisory Committee (09.11). All individuals gave informed consent and/or assent to participate in the study.

### Antigen coupling to magnetic beads

Five different protein constructs of Pfs230 and Pvs230 were included in the Luminex antibody detection assay. Of these, four were from different 6-cysteine domains of Pfs230, namely D1D2, D5D6, D11D12, and D13D14, and the other was from D13 and D14 of Pvs230, assigned as PvD13D14. These constructs were coupled to magnetic Bio-Plex beads (Bio-Rad) following the manufacturer’s instructions and as per prior publications^60^. Briefly, the protein constructs were coupled to uniquely fluorescent microspheres. 50 µL of each microsphere was taken from the stock into a 1.5 mL tube after sonication and vortexing for 15 s and 10 s, respectively. The tubes were immobilized in a magnetic separator rack for 30 – 60 s. The supernatant was discarded, and the remaining microspheres washed with 100 µL of MilliQ water. The suspension was vortexed for 20 s and the supernatant was discarded. The COOH functional group of the microspheres were activated with the addition of 80 µL of monobasic sodium phosphate (100 mM, pH 6.2), 10 µL of sulfo-N-hydrosuccinimide (50 mg/mL) and 10 µL of N-ethyl-N-(3-dimethylaminopropyl) cabodiimide (EDC) (50 mg/mL). The mixture was incubated for 20 min in the dark at RT on a rotator. The microspheres were washed twice with 500 µL of 1X PBS (pH 7.4) and vortexed for 20 s. The activated microspheres were resuspended with 248 µL of 1X PBS, followed by the addition of 2 µg of the antigen. The mixture was then incubated at RT for 2 h on the rotator. Afterwards, the mixture was washed three times with 500 µL of 1X PBS-TBN (PBS, 0.1% (w/v) bovine serum albumin (BSA), 0.02% (v/v) TWEEN-20, 0.05% azide, pH 7) and then resuspended in 125 µL of 1X PBS-TBN. The mixture was kept at 4 °C in the dark until use.

### Multiplex antibody assay

Plasma samples were diluted in PBT (1X PBS, 1% BSA, 0.05% Tween-20) at a dilution of 1:100. Plasma samples from hyper-immune individuals from PNG were used as a positive control pool from which a two-fold serial dilution curve was derived, as previously described^60^. Plasma samples from malaria-naïve individuals from the VBDR were used as the negative control pool. 50 µL of the 1:100 diluted sample was added to a black flat-bottom 96-well plate. A bead mixture with all the coupled microspheres was prepared and 50 µL of this mixture was added to the plate, which was then incubated for 30 min in the dark on a plate shaker. The plate was washed three times with PBST (1X PBS, 0.05% TWEEN-20) using a plate washer. 100 µL of phycoerythrin (PE)-conjugated anti-human secondary antibody (Jackson Immunoresearch) at 1:100 dilution was added to each well and the plate was incubated for 15 min. After a wash, the wells were resuspended with 100 µL PBT and incubated for 5 min on a plate shaker before the plate was read on the MAGPIX instrument. Median fluorescence intensity was obtained and converted to relative antibody units using the PNG serial dilution curve on each plate. This conversion is performed using a five-parameter logistic regression model^60^.

### Production of mRNA-LNP vaccines

#### mRNA production

The mRNAs used in this study were produced using HiScribe T7 mRNA synthesis kit (NEB, Australia) using linearized DNA produced by PCR amplification. The transcribed mRNAs included a 3’-UTR with Kozak sequence, 5’-UTR and polyA_125_ tails. The codon sequences were optimized to reduce the uridine content of mRNA. We used N1-methyl-pseudoUTP instead of UTP to produce chemically modified mRNA, in common with the two approved COVID-19 vaccines. CleanCap reagent AG (TriLink) was used in accordance with the manufacturer’s recommendations to produce Cap1 chemistry at the 5’ terminus. The mRNA was subject to cellulose purification before use.

#### Lipid nanoparticle (LNP) formulation

The following lipids were used in the study: ionisable lipid [(4-hydroxybutyl)-azanediyl]-di-(hexane-6,1-diyl)-bis-(2-hexyldecanoate) (ALC-0315), cholesterol (Sigma-Aldrich, Germany), 1,2-distearoyl-sn-glycero-3-phosphocholine (DSPC) (Avanti Polar Lipids Inc., USA), and the PEGylated lipid methoxypolyethyleneglycoloxy(2000)-N,N-ditetradecylacetamide (ALC-0159). The lipids were used in the proportions used in the BioNTech/Pfizer SARS-CoV-2 mRNA vaccine (*Comirnaty*). Formulations of the vaccines into LNPs involved the following steps: an aqueous solution of mRNA at pH4 was mixed with a solution of the four lipids in ethanol, using a microfluidics mixing device (NxGen Ignite Nanoassemblr) supplied by Precision Nanosystems. The suspension of nanoparticles was adjusted to pH of 7.4 using a 1:3 dilution in Tris buffer, then dialyzed against 25 mM Tris buffer to remove the ethanol. The LNP suspension was adjusted with sucrose solution to produce the cryoprotected, isotonic final form of the product. The product was sterile filtered (0.22 μm) prior to being aliquoted into sterile vials for storage at −80°C. Characterization of the LNPs included analysis for RNA content, encapsulation efficiency and RNA integrity. Particle size and polydispersity index (PDI) were determined by dynamic light scattering, a standard method for submicron dispersions, using a Zetasizer (Malvern Instruments). Typically, encapsulation efficiency was 85-95%, particle sizes (Z-average) of the LNPs were 78 – 81 nm after thawing with PDI < 0.2. All the procedures were carried out by mRNA Core (mrnacore.org).

### Pfs230 D13D14 mRNA-LNPs immunization

Mouse immunization and serum isolation were performed by the WEHI Antibody Facility. Groups of 8–12 week old female C57BL/6 mice (n=6) were immunized with 10 μg of either 230D13D14L or 230D13D14S mRNA-LNP alone. The mRNA-LNPs were administered intramuscularly in a 50 μL dose per mouse. Mice were immunized on day 1 and day 28. For pre-immune sera, mice were bled on day 1 prior to immunization. Mice were also bled on day 21 and day 68 for the first and second bleed, respectively. This work has been approved by the WEHI Animal Ethics Committee 2023.012.

### ELISA for antibody levels

96-well flat-bottomed MaxiSorp plates were coated with 50 μL of 1 μg/mL recombinant Pfs230 D13D14 in 1x PBS for 1 h at RT. All washes were done three times using PBST and all incubations were performed for 1 h at RT. Coated plates were washed and blocked with 1% BSA in PBST for 1 h. Plates were washed and incubated with two-fold serial dilutions of mouse sera starting at 1:200 in 1% BSA PBST, washed as above and incubated with horseradish peroxidase (HRP)-conjugated goat anti-mouse secondary antibody (1:2000; Merck AP124P). 50 μL of ABTS liquid substrate (Sigma 11684302001) was added and incubated at RT and 50 μL of 1% SDS was used to stop the reaction. Absorbance was read at OD 405 nm and all samples were in duplicate. The serial dilution curves were fitted to a third order polynomial regression curve using GraphPad Prism version 10 for Mac (GraphPad Software). End point titers were determined as the reciprocal of the dilution at which the curve intersected the cutoff, and the cutoff was defined as three times the average absorbance values of the respective pre-immune serum.

### Quantification and statistical analysis

Analyses were performed using GraphPad Prism version 10 for Mac (GraphPad Software). Tests and statistics are described in Figure Legends. For end point titers, the serial dilution curves were fitted to a third order polynomial regression curve using GraphPad Prism version 10 for Mac (GraphPad Software). End point titers were determined as the reciprocal of the dilution at which the curve intersected the cutoff, and the cutoff was defined as three times the average absorbance values of the respective pre-immune serum. Automated quantitation of SIFA of macrogametes were performed as follows. Cell segmentation was performed via a custom trained cellpose^94,95^ network on maximum intensity projected brightfield images. Cell scoring was performed via a custom FIJI^92^ macro with manual thresholds set for mean intensity of maximum intensity projected images of 647 and 555 channels. Mask images showing the segmentation and scoring of each cell were produced for manual validation.

## Data and code availability

Cryo-EM maps have been deposited in the EM Data Bank with the following accession codes: EMD-48672 (consensus), EMD-48669 and EMD-48670 (local refinements), and EMD-48673 (composite map). The corresponding atomic coordinates have been deposited in the Protein Data Bank with the accession codes 9MVV (Pfs230 D13D14-Pfs48/45) and 9MVT (Pfs230 D1-D8). Crystal structures have been deposited in the PDB under accession codes 9E7N (Pfs230 D13D14-W2809), 9E7O (Pfs230 D13D14-W2810) and 9E7P (Pfs230 D13D14-W2812). The SDA crosslinking mass spectrometry data have been deposited to the ProteomeXchange Consortium via jPOST^111^ with accession number JPST003585. The mass spectrometry proteomics data have been deposited to the ProteomeXchange Consortium via the PRIDE partner repository^112^ with the dataset identifier PXD060667. The NGS data has been deposited to the European Nucleotide Archive (ENA) under accession number PRJEB85614.

## Acknowledgements

We thank Geoffrey Kong from the Monash Macromolecular Crystallisation Facility (MMCF, Clayton, VIC, Australia) for assistance with setting up the crystallization screens. This research was undertaken using the MX2 beamline at the Australian Synchrotron, part of ANSTO, and made use of the Australian Cancer Research Foundation (ACRF) detector. The authors acknowledge the use of Bio21 Advanced Microscopy Facility, WEHI CryoEM Facility and we would like to thank Andrew Leis for his support with cryo-EM sample preparation and data collection. This work was supported by the MASSIVE HPC facility (www.massive.org.au). NF54/iGP2 was kindly provided by Till Voss (Swiss Tropical and Public Health Institute). We thank Nick Walker and Pailene Lim (WEHI) for assistance with multiplex assays and sample handling. We acknowledge Professor Leanne Robinson (Burnet Institute), Benson Kiniboro and Dr Inoni Betuela (PNGIMR), and the PNGIMR Maprik field team for collection of the human study samples. W-H.T. is supported by National Health and Medical Research Council of Australia (NHMRC) GNT2016908 and APP2001385. Q.G is supported by NHMRC Investigator Grant GNT2007996. R.J.L is supported by NHMRC Investigator Grant GNT1173210. A.G. is a CSL Centenary Fellow. S.S. is supported by the NHMRC Investigator grant (GNT 2016827). C.W.P is supported by Medical Research Future Fund grant MRFCT100006 which funds mrncore.org. The authors acknowledge the Victorian State Government Operational Infrastructure Support and Australian Government NHMRC IRIISS.

## Author contributions

Conceptualization, M.H.D and W-H.T.; Methodology, M.H.D., L-J.C., J.C., K.Z., Q.G., S.F. and R.S.M; Formal analysis, M.H.D., J.C., L-J.C., L.L.T., A.A., T.A.B., L.F.D., R.R., A.A., R.J.L., A.M.V., K.Z., Q.G., A.G., S.S. and W-H.T.; Investigation M.H.D., J.C., L-J.C., L.L.T., A.A., F.M.T.L., M.G, S.L., T.A.B., L.F.D., R.R., A.A., R.M., R.J. L., L.P., P.G. and M.K.; Resources, R.J.L. and I.M.; Writing – Original draft, M.H.D., J.C., L-J.C., L.L.T., A.A., M.T.L., T.A.B., L.F.D. R.R. and R.J.L.; Writing – reviewing and editing, all authors, Visualization, M.H.D., J.C., L-J.C., L.L.T., A.A., F.M.T.L., R.R., A.A., R.J.L., K.Z. and W-H.T.; Supervision, W-H.T, S.L., C.W.P. and M.H.D.; Funding acquisition, Q.C., R.J.L., C.W.P. and W-H.T.

## Materials and Correspondence

Correspondence and requests for materials should be addressed to Wai-Hong Tham. Supplementary Information is available for this paper.

