## Supplemental Figures and Tables for "Cryo-EM structure of endogenous *Plasmodium falciparum* Pfs230 and Pfs48/45 fertilization complex"

FIGURE S1

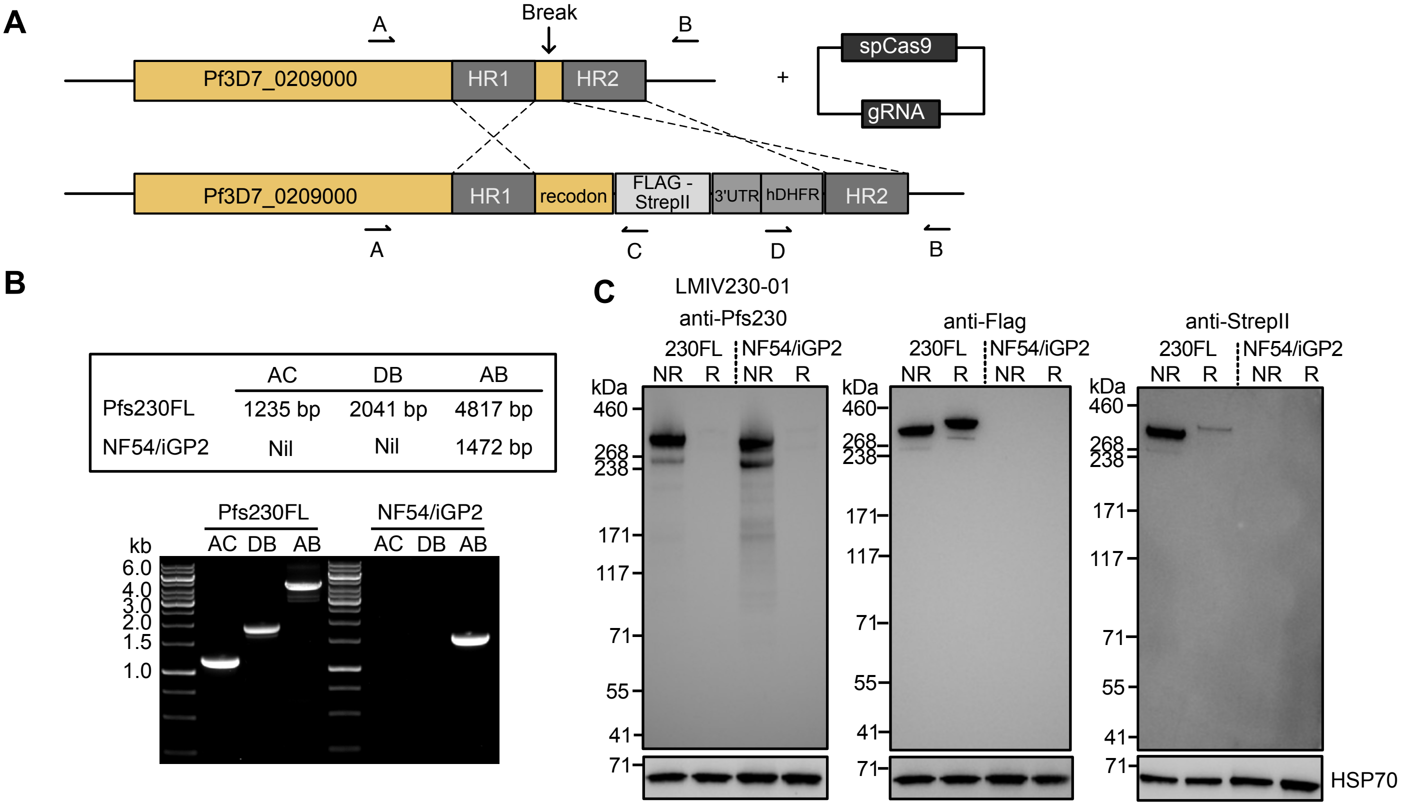

**Figure S1. Transgenic *P. falciparum* line expressing Pfs230 with a C-terminal 3xFLAG-TwinStrepII tag** (A) Schematic illustrating the strategy for generation of the Pfs230-3xFLAG-TwinStrepII line in an NF54/iGP2 background. This transgenic line is referred to as Pfs230FL. Genotyping primers (A, B, C and D) are indicated in the schematic. gRNA, guide RNA; HR, homology region; hDHFR, human dihydrofolate reductase; recodon, recodonized; SpCas9, *Streptococcus pyogenes* Cas9; UTR, untranslated region. 5' and 3' homology regions are indicated as HR1 and HR2, respectively. (B) Expected amplicon sizes are shown in relation to the respective genotyping primer pairs. Gel electrophoresis of the PCR amplicons is shown for both the transgenic Pfs230FL and the parental NF54/iGP2 lines. DNA ladder sizes are shown on the left. (C) Western blot of transgenic Pfs230FL and the parental line NF54/iGP2 using anti-Pfs230 LMIV230-01, anti-FLAG and anti-StrepII antibodies, showing expression of the 3xFLAG and TwinStrepII tag on Pfs230. Pfs230 is present as 360 kDa and 310 kDa bands, with the lower band representing proteolytically cleaved Pfs230. Anti-HSP70 was used as a loading control. Non-reducing and reducing conditions are shown as NR and R, respectively. Molecular weight markers are shown on the left.

**FIGURE S2**

**A**

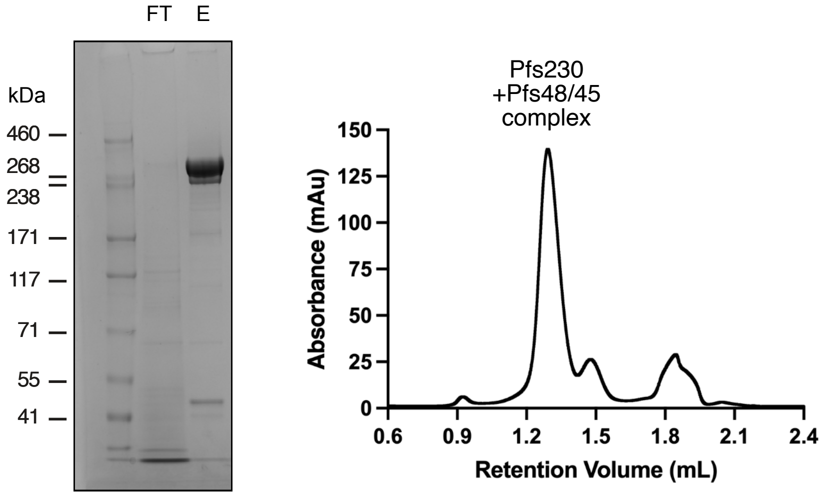

| Protein IDs | Gene names | Mol. weight [kDa] | Peptides | Unique peptides | Sequence coverage % | Intensity |
|---|---|---|---|---|---|---|
| A0A143ZY09 | Pfs230 | 363.2 | 424 | 424 | 75.5 | 774380000 |
| A0A5K1K917;Q8I6T1 | Pfs45/48 | 51.5 | 20 | 20 | 39.1 | 4979700 |
| Q8I2X4 | HSP70 | 72.4 | 26 | 26 | 49.2 | 1585800 |

**B**

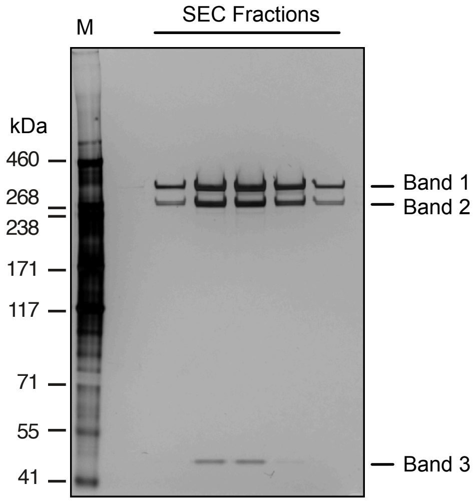

| Band | Gene names | Mol. weight [kDa] | Peptide counts |
|---|---|---|---|
| Band 1 | Pfs230 | 363.2 | 192 |
| Band 2 | Pfs230 | 363.2 | 192 |
| Band 3 | Pfs48/45 | 51.5 | 6 |

**Figure S2. Mass spectrometry analyses confirm purification of endogenous Pfs230-Pfs48/45 fertilization complex**
(A) Coomassie-stained SDS-PAGE gel showing the purification of endogenous Pfs230-Pfs48/45 complex from Pfs230FL using StrepTactin XT resin with the flowthrough (FT) and elution (E) shown. Right panel shows the SEC fractions that correspond with the Pfs230-Pfs48/45 complex. The mass spectrometry analyses show the top three hits with more than three unique peptides from in solution digest preparation. (B) Silver-stained SDS-PAGE gel showing the purification of endogenous Pfs230-Pfs48/45 complex from a Pfs230-HA tagged line using anti-HA, with the relevant SEC fractions shown. Right panel shows the mass spectrometry analyses of the top hit from each band from in gel digest preparation.

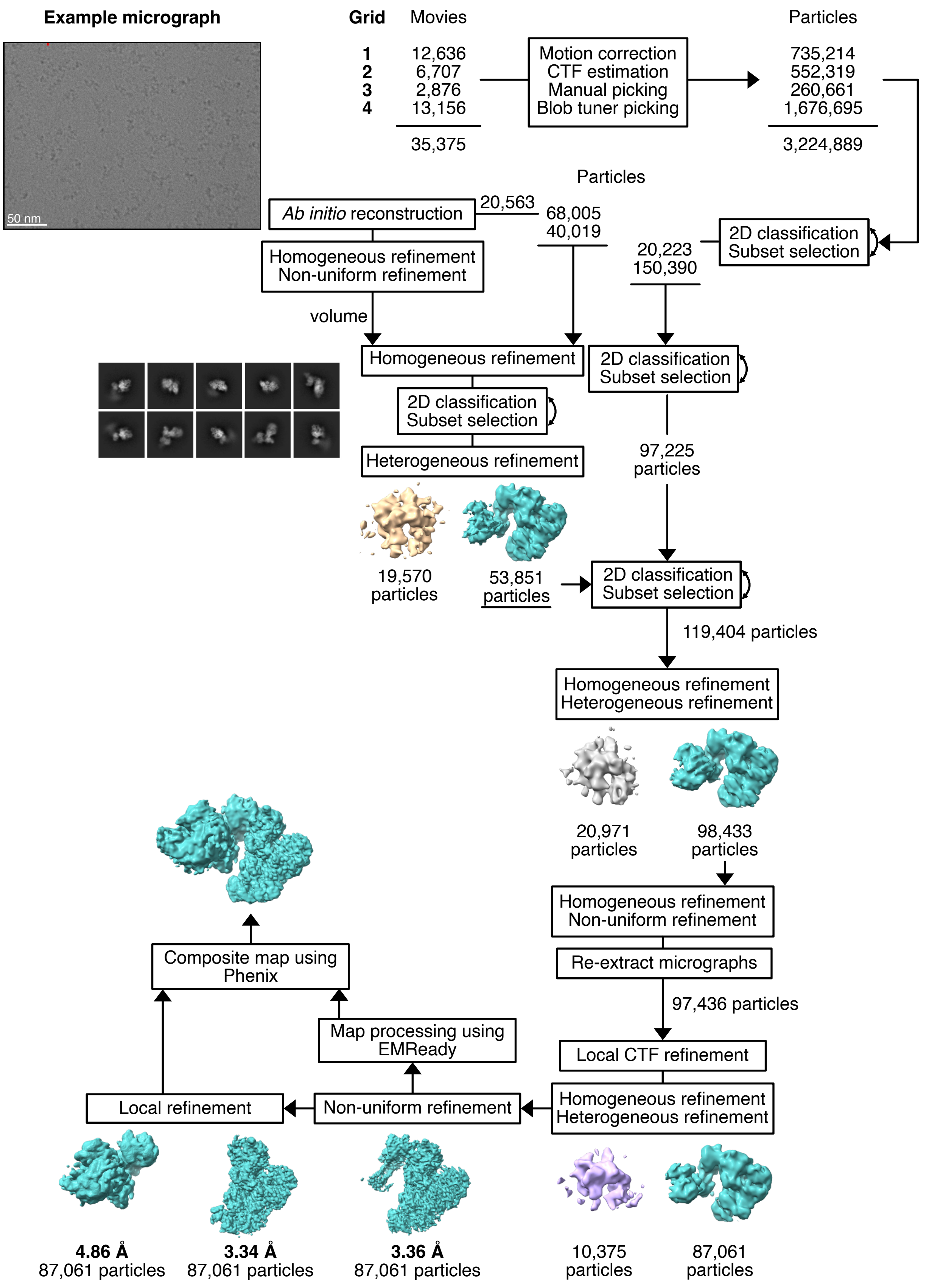

**Figure S3. Workflow for cryo-EM data processing**
Movies from four grids of Pfs230-Pfs48/45 complex after size exclusion chromatography were pre-processed separately in cryoSPARC v4.6.0. Manual particle picking was followed by Blob Tuner Picking. After 2D classification, particles of one grid were used for *ab initio* reconstruction. Particle stacks of the other grids were added at the 2D classification step and further refined using heterogeneous refinement and local contrast transfer function (CTF) refinement. A final non-uniform refinement was used to obtain the consensus map of the complex. The map was further locally refined to improve the map of the lower resolution part of Pfs230 (domains 1-8) and the higher resolution part containing Pfs230 domains 9-14 and Pfs48/45 using soft masks around the regions of interest. EMReady (v2.0) was used to improve the quality of these maps further. A composite map was generated from these maps using Phenix (v1.21.1).

**FIGURE S4**

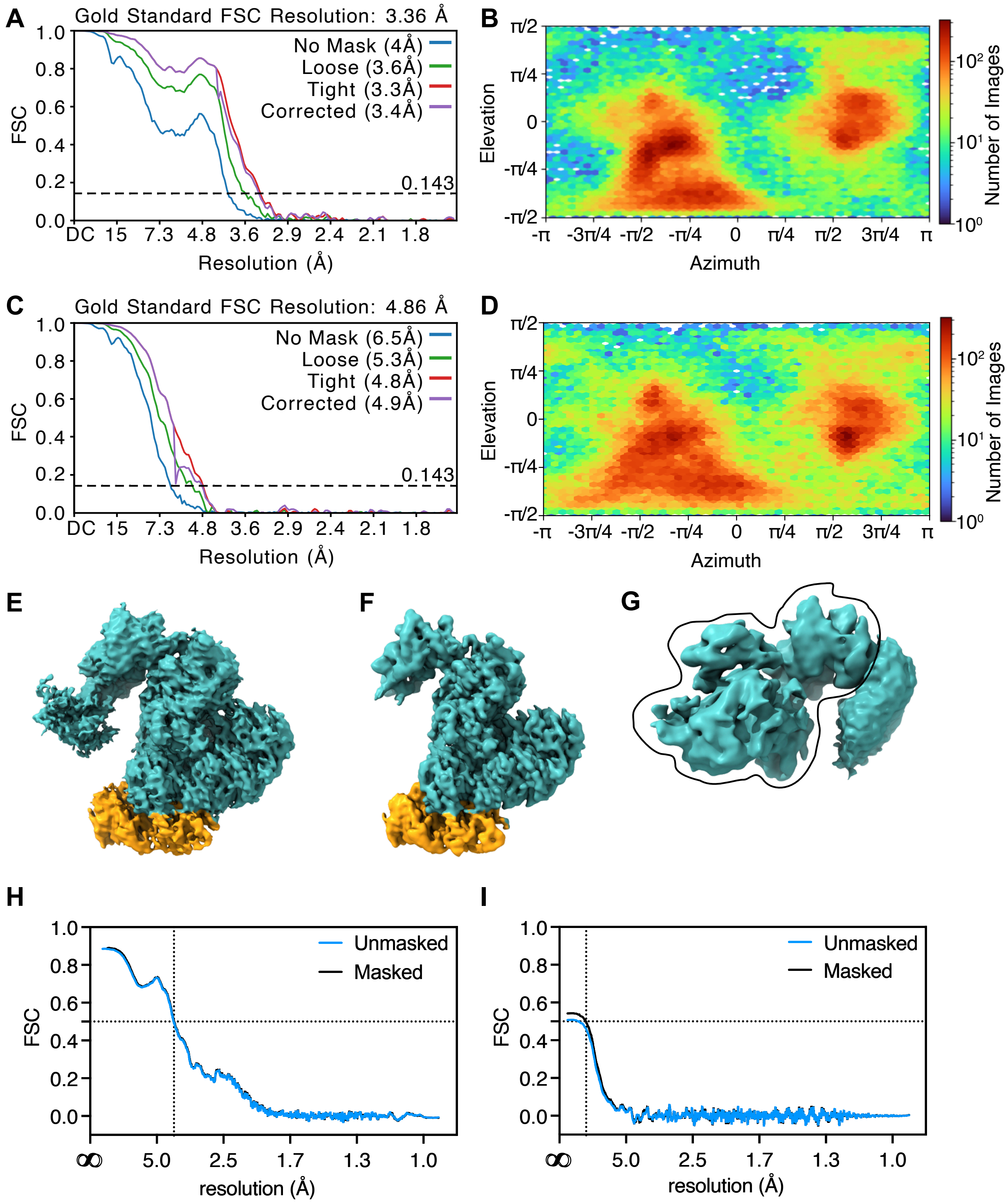

**Figure S4. Cryo-EM processing statistics**

(A) Gold-Standard Fourier shell correlation (FSC) curves of the consensus Pfs230-Pfs48/45 reconstruction. (B) Particle view distribution of the consensus Pfs230-Pfs48/45 reconstruction. (C) Gold-Standard FSC curves of the local refinement map corresponding to Pfs230 D1 to D8 reconstruction. (D) Particle view distribution of the local refinement reconstruction. (E) Consensus map after non-uniform refinement. (F) Consensus map after post-processing with EMReady (v2.0). (G) Local refinement map of the Pfs230 D1 to D8 region. Surrounding trace indicates mask used for local refinement. Particle stacks were not subtracted. (H) Model vs map FSC curves of Pfs230 D9 to D14-Pfs48/45 model. (I) Model vs map FSC curves for model comprising Pfs230 D1 to D8.

**FIGURE S5**

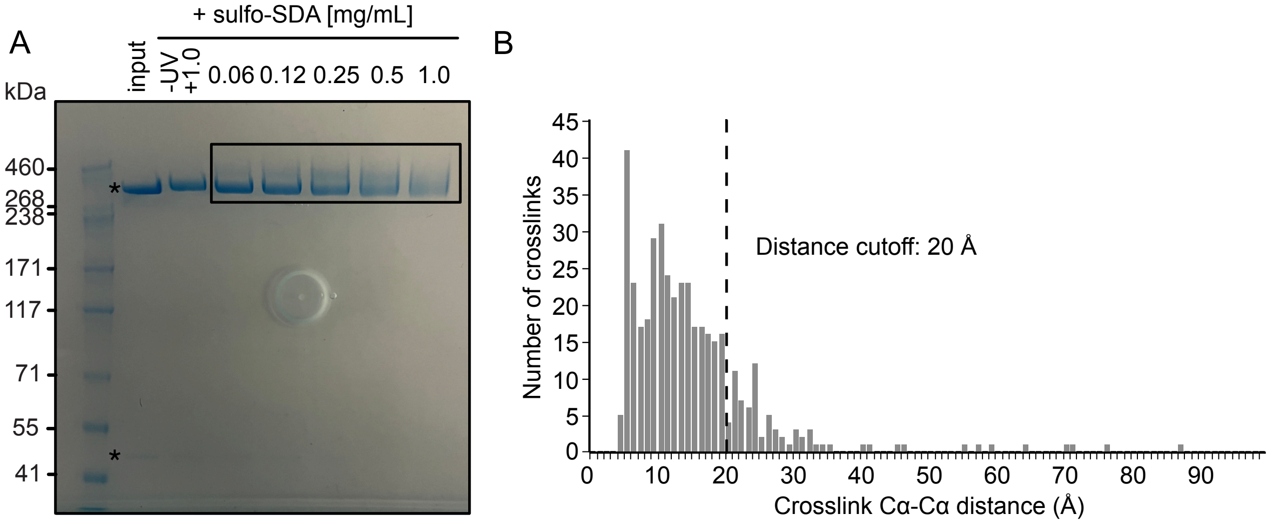

**Figure S5. Crosslinking mass spectrometry of the Pfs230-Pfs48/45 complex**
(A) SDS-PAGE gel of purified Pfs230-Pfs48/45 complex without crosslinker (input), and after incubation with crosslinker Sulfo-SDA at different concentrations and UV exposure. Pfs230 and Pfs48/45 bands are indicated (*) at 360 and 50 kDa, respectively, in the input sample. Covalent linkage of the two proteins results in protein bands at a higher molecular weight. Rectangular box indicates bands used for mass spectrometry. (B) Histogram showing the distance distribution of crosslinks using the structural model of Pfs230-Pfs48/45. The line at 20 Å indicates the distance cutoff for links classified as long-distance. 82% of crosslinks fall within the crosslinker specific distance cutoff.

**FIGURE S6**

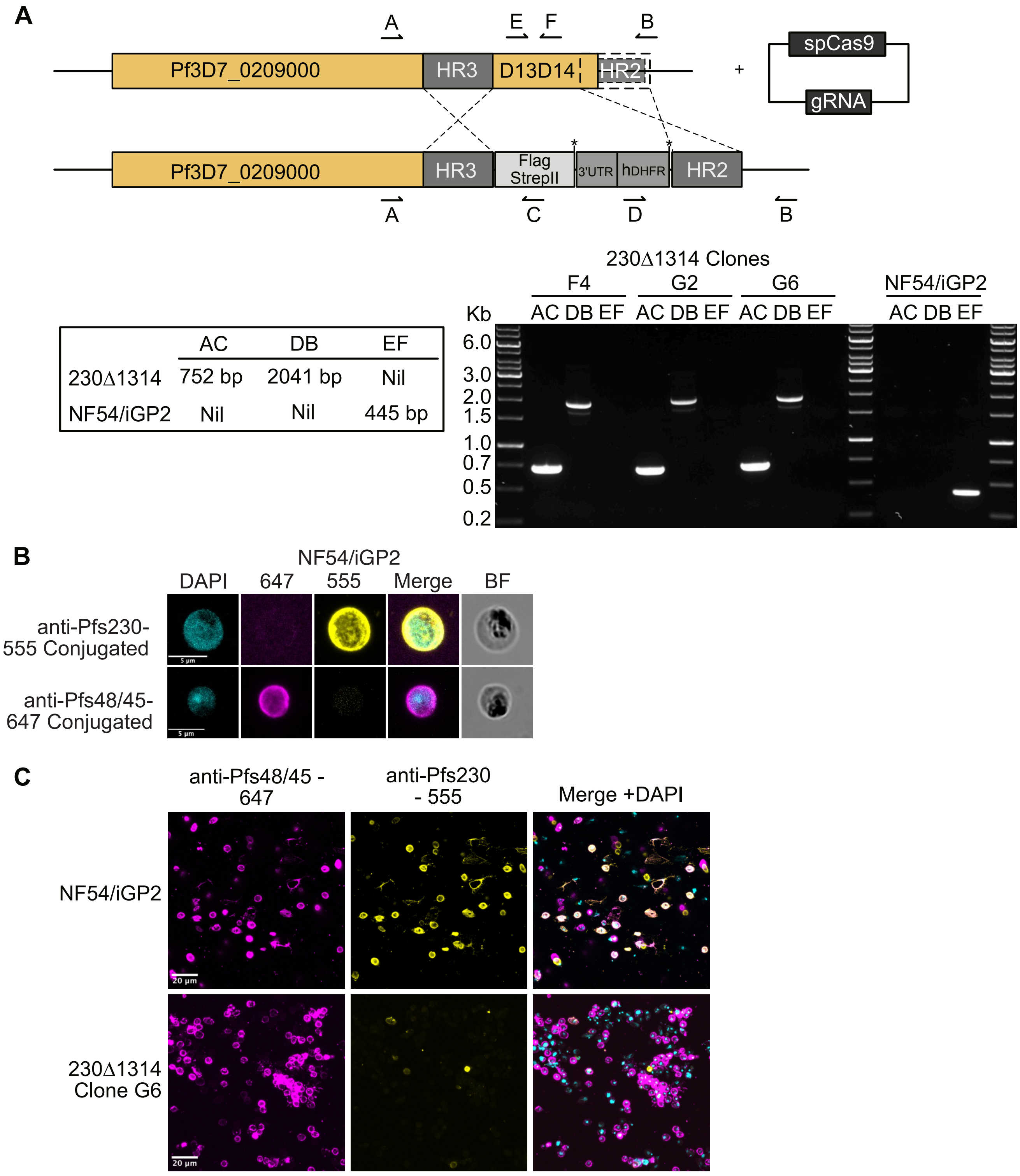

**Figure S6. Generation of transgenic 230Δ1314 and localization of Pfs230 in activated macrogametes** (A) Schematic illustrating the strategy for generation of transgenic 230Δ1314 with 3xFLAG-TwinStrepII in NF54/iGP2 background. Genotyping primers are indicated in the schematic. Expected amplicon sizes are shown in relation to the respective genotyping primer pairs (bottom panel, left). Gel of the PCR amplicons is shown for both the transgenic 230Δ1314 and the parental line NF54/iGP2 lines (bottom panel, right). DNA ladder sizes are shown on the left. (B) Representative surface immunofluorescence assay (SIFA) images of NF54/iGP2 macrogametes which express full-length Pfs230 stained individually with anti-Pfs48/45 conjugated to 647 and anti-Pfs230 conjugated to 555 show no cross-over fluorescence. DAPI, merge and bright field (BF) images are shown. Scale bar = 5 µm. (C) Representative full field of view images taken for quantitation of SIFA of activated macrogametes in the transgenic 230Δ1314 and the parental line NF54/iGP2 lines stained with both anti-Pfs48/45 conjugated to 647 and anti-Pfs230 conjugated to 555. DAPI and merged images are shown. Scale bar = 20 µm.

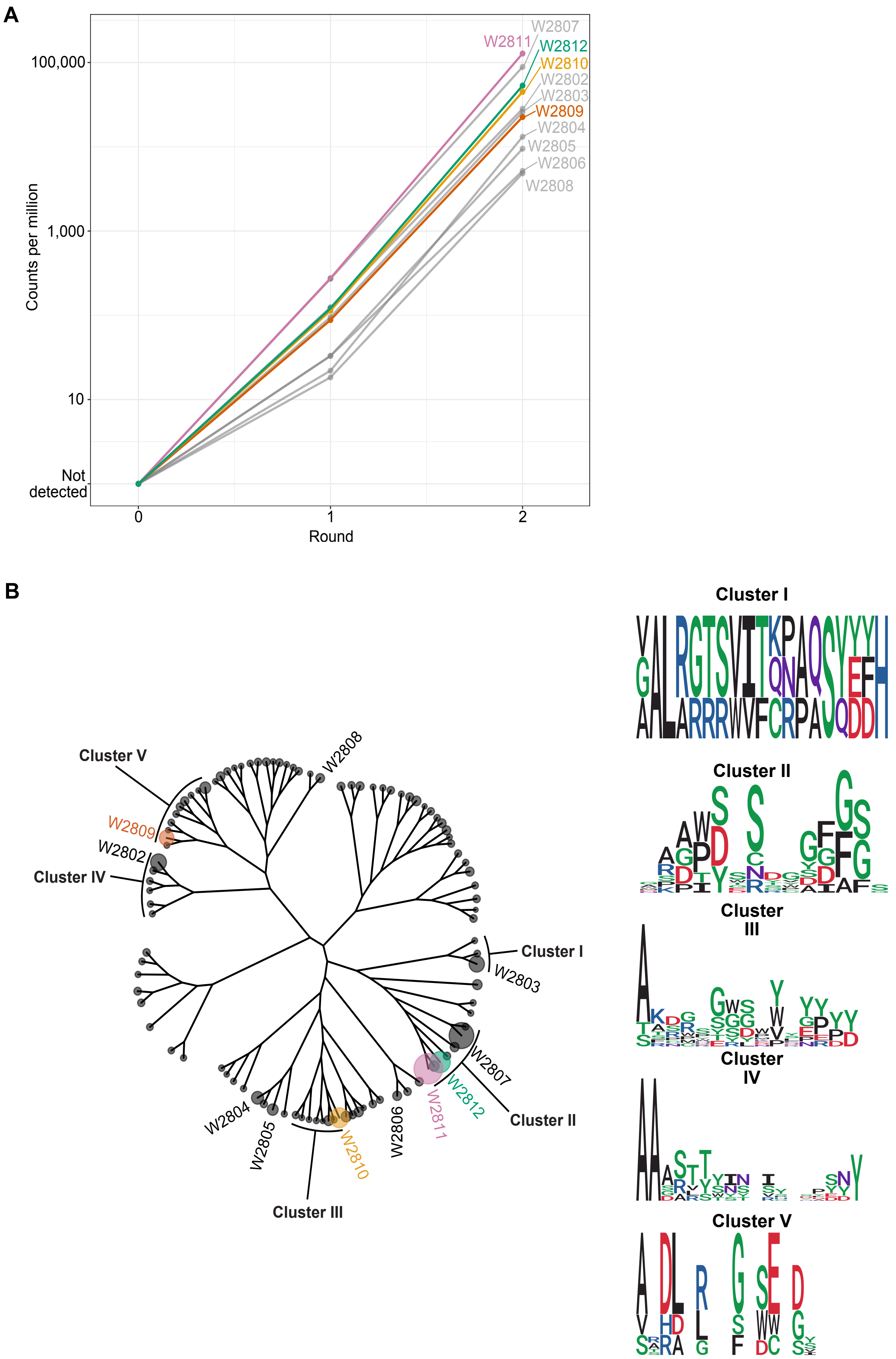

**Figure S7. Next-Generation Sequencing analyses of Pfs230 D13D4 nanobodies phage display selection rounds.** (A) Line graph of the normalized counts per million (CPM) of Pfs230 D13D14 nanobodies across the panning process. (B) Cladogram of the 100 most abundant nanobodies from Next-Generation Sequencing (NGS) after two rounds of phage display. Anti-Pfs230 nanobodies identified by Sanger sequencing are labelled. Tree tips are scaled relative to abundance (counts per million). Selected nanobody clusters (I to V) are highlighted and amino acid sequences of CDR3s in selected clusters of the cladogram are visualized as sequence logos.

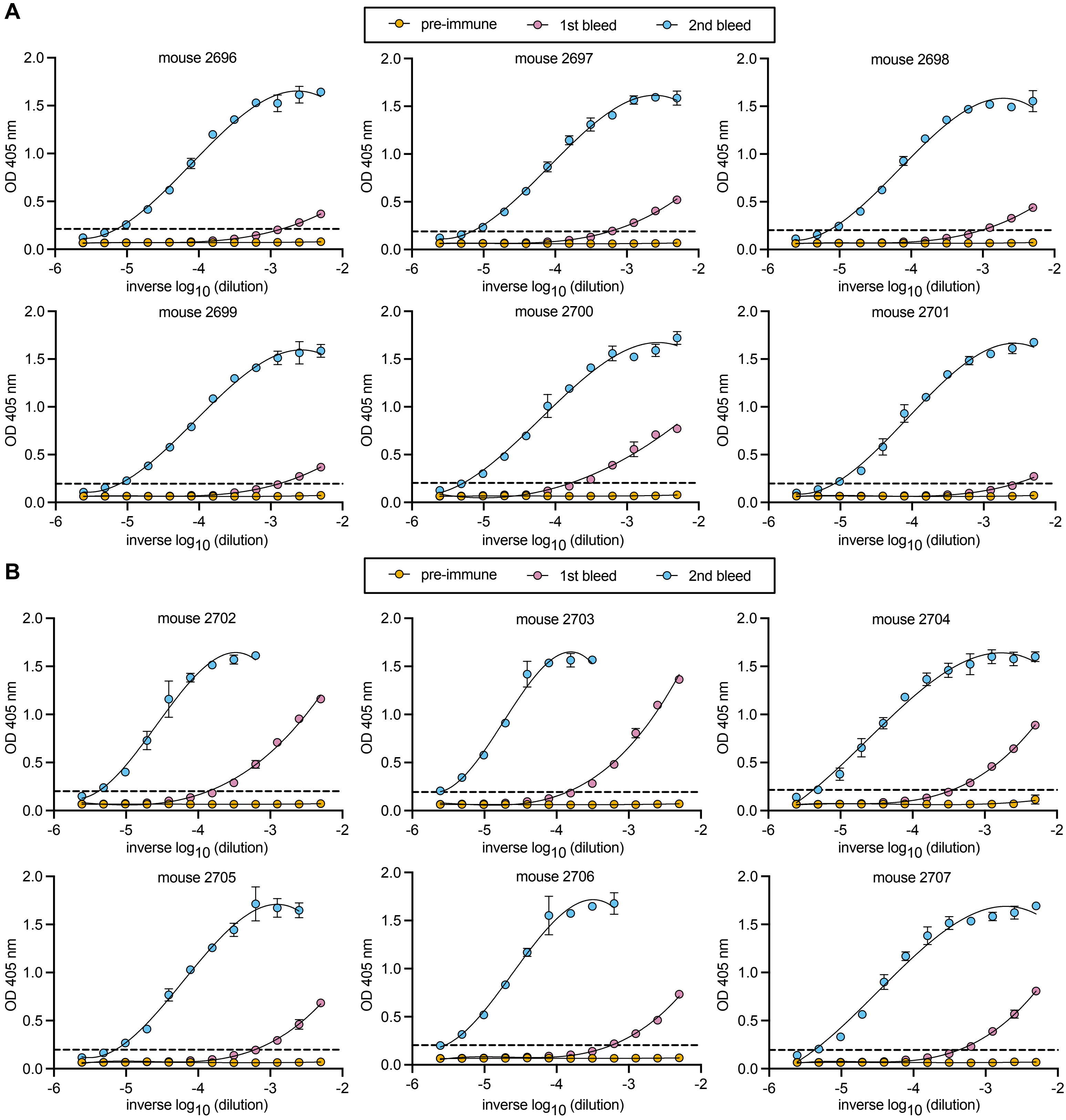

**Figure S8. Immunization with Pfs230 D13D14 mRNA-LNPs elicited antibodies against Pfs230 D13D14.**

(A) Sera of mice immunized with Pfs230 D13D14L mRNA-LNP and (B) Pfs230 D13D14S mRNA-LNP were serially diluted two-fold and binding to Pfs230 D13D14 was measured by ELISA. Serum dilution series was performed on pre-immune (orange), first bleed (pink) and second bleed (blue) sera for each mouse. Curves were fitted to a third order polynomial regression curve and outliers were excluded. The cutoff to determine endpoint titer is represented by the dashed line and was defined as three times the average absorbance values of the respective mouse pre-immune serum. Error bars represent standard deviation of the mean of two technical replicates.

Table S1. Cryo-EM data collection, refinement and validation statistics.

| | Pfs230-Pfs48/45 complex | | |
|---|---|---|---|
| | Consensus map | Local refinement – Pfs230 D1-D8 | Local refinement – Pfs230 D9-14 and Pfs48/45 |
| EMD-, PDB | EMD-48672, 9MVV | EMD-48669, 9MVT | EMD-48670 |
| | EMD-48673 (composite EM map) | | |
| **Data collection and processing** | | | |
| Microscope | Titan Krios | | |
| Detector | Falcon 4 | | |
| Magnification | 96,000 | | |
| Voltage (kV) | 300 | | |
| Electron exposure ($e^-/Å^2$) | 50 | | |
| Defocus range (μm) | -0.4 to -1.6 | | |
| Pixel size (Å) | 0.808 | | |
| Number of micrographs | 35,375 | | |
| Symmetry imposed | C1 | C1 | C1 |
| Initial particle images (no.)[a] | 3,224,889 | 3,224,889 | 3,224,889 |
| Final particle images (no.) | 87,061 | 87,061 | 87,061 |
| Map resolution (Å) | 3.36 | 4.86 | 3.34 |
| FSC threshold | 0.143 | 0.143 | 0.143 |
| Map resolution range (Å) | 2.9-50.9 | | |
| **Refinement** | [b] | | |
| Initial model used | AlphaFold2, 7ZXG, 7ZXF | AlphaFold2 | |
| Model resolution (Å) | 3.98 | 12.46 | |
| FSC threshold | 0.5 | 0.5 | |
| Model composition | Pfs230 D9-D14 and Pfs48/45 | Pfs230 D1-D8 | |
| Nonhydrogen atoms | 10436 | 7304 | |
| Protein residues | 1394 | 1471 | |
| **B factors ($Å^2$)** | | | |
| Protein | 30.0/ 293.7/ 104.3 | 30.0/ 791.4/ 395.9 | |
| R.m.s. deviations | | | |
| Bond lengths (Å) | 0.004 | 0.003 | |
| Bond angles (°) | 0.767 | 0.842 | |
| **Validation** | | | |
| Ramachandran plot | | | |
| outliers (%) | 0.0 | 0.14 | |
| favored (%) | 93.59 | 93.52 | |
| allowed (%) | 6.41 | 6.34 | |
| Rotamer outliers (%) | 0.0 | 0.0 | |
| Cβ outliers (%) | 0.0 | NA | |
| MolProbity score | 1.74 | 1.73 | |
| Clashscore | 5.89 | 5.61 | |
| **Model vs. data fit** | | | |
| CC (mask) | 0.71 | 0.53 | |
| CC (box) | 0.86 | 0.70 | |

a: Blob tuner picked particles, b: EMReady processed map. CC, correlation coefficient; EMD-, Electron Microscopy Data (accession code); FSC, Fourier shell correlation; PDB, Protein Data Bank (accession code); R.m.s, root mean square.

Table S2. Data collection and refinement statistics for Pfs230 D13D14-nanobody complexes.

| | **Pfs230 D13D14 - W2809** | **Pfs230 D13D14 – W2810** | **Pfs230 D13D14 – W2812** |
|---|---|---|---|
| PDB ID | 9E7N | 9E7O | 9E7P |
| **Data collection statistics** | | | |
| Wavelength (Å) | 0.953732 | 0.953647 | 0.953728 |
| Space group | $P4_3\ 2_1\ 2$ | C2 | $P4_12_12$ |
| Cell axes (Å) (a, b, c) | 79.9, 79.9, 342.7 | 135.5, 62.8, 56.5 | 134.4, 134.4, 75.6 |
| Cell angles (°) (α, γ, β) | 90, 90, 90 | 90, 98.9, 90 | 90, 90, 90 |
| Resolution range (Å) | 47.17-2.49 (2.64-2.49) | 39.92-1.93 (2.05-1.93) | 47.60-3.22 (3.41-3.22) |
| Completeness (%) | 99.8 (98.6) | 99.5 (98.5) | 99.9 (99.5) |
| Total no. of reflections | 1,062,663 (165,943) | 247601 (38089) | 156973 (24880) |
| Unique reflections | 73732 (11916) | 69078 (10991) | 21640 (3479) |
| Redundancy | 14.4 (13.9) | 3.6 (3.5) | 7.2 (7.2) |
| $R_{meas}$ (%) | 16.5 (232.7) | 8.8 (137.1) | 13.1 (147.2) |
| $CC_{1/2}$ (%) | 99.9 (53.1) | 99.8 (52.5) | 99.9 (54.7) |
| I/σ | 11.75 (1.09) | 9.95 (1.06) | 12.13 (1.30) |
| Wilson B ($Å^2$) | 68.3 | 43.97 | 95.58 |
| **Refinement statistics** | | | |
| $R_{work}/R_{free}$ (%) | 22.4 / 26.4 | 21.4 / 25.4 | 28.6 / 32.0 |
| No. of atoms | | | |
| Protein | 6686 | 3206 | 3147 |
| Water | 62 | 128 | |
| B factors ($Å^2$) | | | |
| Chain A | 66.3 | 51.7 | 112.6 |
| Chain B | 59.8 | 49.8 | 110.9 |
| Chain C | 74.9 | | |
| Chain D | 76.4 | | |
| Water | 56.4 | 45.9 | |
| R.m.s. deviations | | | |
| Bond lengths (Å) | 0.004 | 0.009 | 0.003 |
| Bond angles (°) | 0.672 | 1.028 | 0.602 |
| Ramachandran plot | | | |
| outliers (%) | 0.0 | 0.0 | 0.0 |
| favored (%) | 96.3 | 97.7 | 94.5 |
| Rotamer outliers (%) | 0.28 | 0.59 | 0.31 |
| C-beta outliers | 0.0 | 0.0 | 0.0 |
| MolProbity score | 1.42 | 1.40 | 1.82 |

For each crystal structure a single crystal was used for data collection.
* The values in parentheses represent the highest-resolution shell.

**Table S3**. Interactions Pfs230 D13D14 – W2809 complex: chain **B-D A2** interface area 623.1 A$^2$ / chain CA interface area 711.3 A$^2$

| Pfs230 | Group | W2809 | Location | Group | Distance |
|---|---|---|---|---|---|
| | | Hydrogen bonds | | | |
| Gln 2978 | OE1 | Arg 30 | CDR1 | NH2 | 2.4 |
| Tyr 2980 | O | Ser 56 | CDR2 | OG | 2.6 |
| Glu 2982 | OE1 | Ser 52 | CDR2 | OG | 2.9 |
| Glu 2982 | OE2 | Ser 54 | CDR2 | N | 3.0 |
| Glu 2982 | OE1 | Ser 54 | CDR2 | OG | 2.6 |
| His 2994 | NE2 | Asp 31 | CDR1 | O | 2.8 |
| Ser 2999 | O | Gly 103 | CDR3 | N | 2.9 |
| Ser 2999 | O | Gly 104 | CDR3 | N | 2.9 |
| Ser 2999 | N | Gly 104 | CDR3 | O | 3.1 |
| | | Salt bridges | | | |
| His 2942 | NE2 | Asp 31 | CDR1 | OD1 | 2.9 |
| Lys 2943 | NZ | Asp 31 | CDR1 | OD1 | 3.3 |
| | | Other interfacing residues in Pfs230 | | | |
| Tyr 2940 | Lys 2981 | His 2984 | Ser 2993 | Phe 2996 | Thr 2997 |
| Tyr 2998 | Lys 3000 | Lys 3001 | Cys 3010 | Ile 3109 | |
| | | Other interfacing residues in W2809 | | | |
| Trp 53 | Ile 57 | Asn 74 | Trp 100 | Pro 101 | Gly 102 |
| Met 1105 | Trp 106 | | | | |

Interactions Pfs230 D13D14 – W2810 complex: chain **AB** interface area 588.0 A$^2$

| Pfs230 | Group | W2810 | Location | Group | Distance |
|---|---|---|---|---|---|
| | | Hydrogen bonds | | | |
| Tyr 2879 | OH | Asp 104 | CDR3 | OD2 | 3.4 |
| Asn 2882 | N | Ser 103 | CDR3 | OG | 3.0 |
| Asn 2882 | ND2 | Ser 102 | CDR3 | OG | 2.9 |
| Asn 2882 | OD1 | Ser 102 | CDR3 | N | 3.1 |
| Asn 2882 | OD1 | Ser 103 | CDR3 | OG | 3.6 |
| Val 2892 | N | Ser 103 | CDR3 | O | 3.1 |
| Phe 2895 | O | Tyr 105 | CDR3 | N | 2.8 |
| | | Salt bridges | | | |
| Lys 2900 | NZ | Glu 44 | CDR1 | OE1 | 3.0 |
| Lys 2900 | NZ | Glu 44 | CDR1 | OE2 | 3.5 |
| Lys 2931 | NZ | Asp 104 | CDR3 | OD2 | 2.9 |
| Lys 2931 | NZ | Asp 104 | CDR3 | OD1 | 3.0 |
| | | Other interfacing residues in Pfs230 | | | |
| Pro 2880 | Thr 2881 | Glu 2883 | Glu 2888 | Asn 2889 | Phe 2890 |
| Phe 2891 | Asn 2896 | Leu 2897 | Asn 2910 | Asp 2922 | Tyr 2924 |
| Lys 2929 | Leu 2955 | | | | |
| | | Other interfacing residues in W2810 | | | |
| Ala 33 | Phe 37 | Phe 47 | Ser 50 | Ser 52 | Gly 56 |
| Ser 57 | Ile 58 | Arg 59 | Pro 100 | Tyr 101 | Gly 106 |
| Phe 108 | | | | | |

Interactions Pfs230 D13D14 – W2812 complex: chain **AB** interface area 685.2 A$^2$

| Pfs230 | Group | W2812 | Location | Group | Distance |
|---|---|---|---|---|---|
| | | Hydrogen bonds | | | |

| | | | | | |
|---|---|---|---|---|---|
| Lys 2858 | N | Asn 105 | CDR3 | OD1 | 3.0 |
| Lys 2858 | O | Ile 104 | CDR3 | N | 3.1 |
| Ser 2954 | O | Ser 57 | CDR2 | OG | 2.9 |
| Lys 2959 | NZ | Tyr 59 | CDR2 | OH | 2.9 |
| Gln 2962 | O | Thr 109 | CDR3 | N | 2.9 |
| Gln 2962 | O | Thr 109 | CDR3 | OG1 | 3.4 |
| Asn 2963 | OD1 | His 106 | CDR3 | ND1 | 2.8 |
| Ile 2964 | O | Ile 107 | CDR3 | N | 3.2 |
| Ile 2964 | N | Ile 107 | CDR3 | O | 2.8 |
| Tyr 2966 | N | Asn 105 | CDR3 | O | 2.4 |
| Tyr 2966 | OH | Ala 58 | CDR2 | O | 3.2 |
| Tyr 2966 | O | Asn 105 | CDR3 | ND2 | 2.7 |
| Other interfacing residues in Pfs230 | | | | | |
| Leu 2897 | Glu 2859 | His 2860 | Leu 2955 | Leu 2957 | Asn 2961 |
| Ile 2965 | Gly 2967 | Asn 2968 | | | |

| Other interfacing residues in W2812 | | | | | |
|---|---|---|---|---|---|
| Leu 47 | Thr 52 | Arg 53 | Ser 54 | Val 56 | Thr 102 |
| Tyr 103 | Tyr 108 | Asn 110 | Ser 112 | Asn 113 | |

Table S4. Primers and synthetic DNA sequences, related to Figure 1 and Figure 3.

| **Primers** | | |
|---|---|---|
| **Name** | **Sequence** | **Amplification product** |
| Pfs230FL 3xFlag-TwinStrepII genotyping | | |
| (**A**) Pfs230 tag 5' int check F | GATGTACCTTCGAAAACTATAACAGC | 5' integration check of tag 1253 bp |
| (**C**) FLAG int check R | TAATCCTTATCATCGTCGTCCTTG | |
| (**D**) hDHFR_1.2KOInt_F | CCGCTCAGGAACGAATTTAG | 3' integration check of tag 2041 bp |
| (**B**) Pfs230 tag 3' int check R | CGAAGATGTGGAAGGACTC | |
| (**A**) Pfs230 tag 5' int check F | GATGTACCTTCGAAAACTATAACAGC | Over region which tag is inserted 1721 bp (wild type) 4817 bp (integrated) |
| (**B**) Pfs230 tag 3' int check R | CGAAGATGTGGAAGGACTC | |
| 230Δ1314-3xFlag-TwinStrepII genotyping | | |
| (**A**) Pfs230 5' int check F2 | AAGTTACTGGAGATGAAACAGCTAC | 5' integration check of D13D14 deletion/tag insert 752 bp |
| (**C**) FLAG int check R | TAATCCTTATCATCGTCGTCCTTG | |
| (**D**) hDHFR_1.2KOInt_F | CCGCTCAGGAACGAATTTAG | 3' integration check of tag 2041 bp |
| (**B**) Pfs230 tag 3' int check R | CGAAGATGTGGAAGGACTC | |
| (**E**) 230-D13D14 WT check F | ATGTTACTACTAAAGTTGCTACTTG | Over the Pfs230 D13D14 coding sequence which is deleted 445 bp |
| (**F**) 230-D13D14 WT check R | CATCCATGAATTTCTTTATATCCTTG | |
| 230Δ1314-3xFlag-TwinStrepII line generation | | |
| 230-D13D14-KO 5'HR F | GGTGCGGCCGCTCAAACAGAAATATTATTCATGG | Amplifies 498 bp of nucleotide sequence directly 5' of the K2828 Pfs230 Domain 13 |
| 230-D13D14-KO_5'HR R | GGTCTCGAGTTTTTCATCTATTTTTATTTTATCCATTGTAC | |
| **Guide RNA target sequence** | | |
| Pfs230_g8947 | ATTCATGGATGTGATTTCAC | |
| **Recodonised Coding Sequence (Pfs230 CDS)** | | |
| **Original sequence:**<br>TCATGGATGTGATTTCACAGGAAAATATTCCCATTTATTTACATATTCAAAAAAACCTTTACCAAATGATGATGATAT<br>ATGTAATGTAACTATAGGTAATAATACATTCTCAGGTTTTGCATGCTTAAGCCATTTTGAATTAAAACCAAATAACTG<br>CTTCTCATCTGTTTATGATTATAATGAAGCCAATAAAGTTAAAAAATTATTCGATCTATCCACAAAAGTAGAATTAGA<br>CCATATCAAACAAAATACTTCAGGATATACACTATCATATATTATTTTTAATAAAGAATCCACAAAACTTAAATTCTC<br>ATGTACATGCTCATCCAACTATTCAAATTATACTATACGAATCACATTTGATCCTAATTATATAATCCCAGAACCTCA<br>ATCAAGAGCCATCATTAAATATGTAGATCTGCAAGATAAAAATTTTGCAAAATACTTGAGAAAGCTT<br>**Recodonised sequence:**<br>CCATGGCTGCGACTTTACTGGAAAATATTCCCACCTGTTCACCTATAGCAAAAAGCCGCTGCCGAACGACGA<br>CGACATCTGCAACGTGACCATTGGCAACAACACCTTCTCCGGTTTTGCATGTCTGTCCCACTTTGAACTCAAG | | |

| |
|---|
| CCAAATAATTGTTTCTCTAGCGTCTATGATTATAACGAGGCTAATAAGGTGAAAAAGCTTTTCGACTTGTCGACCAAAGTTGAGCTGGATCATATTAAGCAGAATACCAGCGGTTATACTCTGAGTTACATCATCTTTAACAAAGAGAGCACCAAGTTGAAGTTCAGCTGCACGTGCAGCAGCAATTACTCTAACTACACGATCCGCATTACCTTCGACCCGAACTACATCATTCCGGAACCGCAGAGCCGTGCGATTATCAAGTACGTTGATCTGCAAGATAAAAACTTTGCGAAATACCTGCGTAAATTG |
| **gBlock DNA Synthesis** |
| T7-primer-adaptors_3xFlag-TwinStrepII (Restriction sites in bold, linker regions unmarked, as well as additional stop codons) |
| TAATACGACTCACTATAGGG**CTCGAG**GCAGCAGCCGACTACAAAGATGACGACGATAAAGATTACAAGGACGACGATGATAAGGATTATAAAGATGATGACGATAAGGCCGCAGCCTCAGCCTGGAGTCATCCACAGTTTGAGAAAGGAGGTGGTAGTGGAGGAGGATCAGGTGGATCTGCCTGGTCTCATCCTCAATTTGAAAAATGATAATAG**CCCGGG**CCGCTGAGCAATAACTAGC |
